# A *Drosophila* model of neuronal ceroid lipofuscinosis *CLN4* reveals a hypermorphic gain of function mechanism

**DOI:** 10.1101/579771

**Authors:** Elliot Imler, Jin Sang Pyon, Selina Kindelay, Yong-quan Zhang, Sreeganga S. Chandra, Konrad E. Zinsmaier

## Abstract

The autosomal dominant neuronal ceroid lipofuscinoses (NCL) *CLN4* is caused by mutations in the synaptic vesicle (SV) protein CSPα, which is a critical co-chaperone of Hsc70 protecting synapses from activity-dependent degeneration. We developed the first animal models of *CLN4* by expressing either *CLN4* mutant human CSPα (hCSPα) or *Drosophila* CSP (dCSP) in fly neurons. Similar to patients, *CLN4* mutations induced excessive oligomerization of mutant hCSPα and premature lethality in a dose-dependent manner. Instead of being localized to SVs, most *CLN4* mutant hCSPα abnormally accumulated in axons and somata, and co-localized with ubiquitinated proteins and the prelysosomal markers HRS and LAMP1. Ultrastructurally, abnormal multi-laminar membrane structures were frequently observed in axons and somata next to degenerative abnormalities. The lethality, oligomerization and prelysosomal accumulation induced by *CLN4* mutations was attenuated by reducing wild type (WT) dCSP levels and enhanced by increasing WT dCSP or hCSPα levels, which indicates that both *CLN4* alleles resemble dominant hypermorphic gain of function mutations. Furthermore, reducing the gene dosage of Hsc70 also attenuated *CLN4* phenotypes. Taken together, we suggest that *CLN4* alleles resemble dominant hypermorphic gain of function mutations that drive excessive oligomerization and impair membrane trafficking.

## Introduction

NCLs comprise a group of progressive neurodegenerative diseases with 14 known disease-associated genes, termed *CLN1-14* (Haltia, 2003; Haltia and Goebel, 2013; Jalanko and Braulke, 2009; Mole and Cotman, 2015). NCLs have mostly an infantile or juvenile symptomatic onset and are characterized by loss of vision, gait abnormalities, seizures, dementia, and premature death. In general, NCLs are considered lysosomal storage diseases due the accumulation of lipofuscin and are typically caused by recessive loss of function mutations with one exception, *CLN4*. Accordingly, lysosomal dysfunction, dysregulated ER-lysosomal trafficking, or aberrant lipid modifications are thought to be the basis for most NCLs, consistent with the known function of the mutated genes (Bennett and Rakheja, 2013; Carcel-Trullols et al., 2015; Mole and Cotman, 2015; Warrier et al., 2013). Understanding disease mechanisms of NCLs has implications as well for more prevalent diseases since various mutations in a growing number of CLN genes cause also other diseases like frontotemporal lobar degeneration, progressive epilepsy with mental retardation, spinocerebellar ataxia, retinitis pigmentosa, juvenile cerebellar ataxia, or Parkinson disease 9 (Bras et al., 2012; Mole and Cotman, 2015; Yu et al., 2010).

The autosomal dominant inherited NCL *CLN4* has an adult onset between 25-46 years. *CLN4* is caused by either the amino acid (aa) substitution L115R or the single amino acid deletion L116Δ in the SV protein CSPα, which is encoded by the human *DNAJC5* gene (Benitez et al., 2011; Cadieux-Dion et al., 2013; Noskova et al., 2011; Velinov et al., 2012). CSPα is unique among NCL-associated genes since it encodes a SV protein with no known lysosome-associated functions. Accordingly, there is no *CLN4* model explaining lysosomal failure.

CSPα is an evolutionary conserved neuroprotective co-chaperone of Hsc70 and required to maintain synaptic function and prevent neurodegeneration (Burgoyne and Morgan, 2011; Burgoyne and Morgan, 2015; Zinsmaier, 2010). Gene deletions in flies and mice cause progressive locomotor defects, paralysis and premature death due to neurodegeneration (Chandra et al., 2005; Fernandez-Chacon et al., 2004; Umbach et al., 1994; Zinsmaier, 2010; Zinsmaier et al., 1994). On SVs, CSPα forms a molecular chaperone complex with Hsc70 for a selected set of clients, which include SNARE proteins and dynamin (Chandra et al., 2005; Nie et al., 1999; Sharma et al., 2012a; Sharma et al., 2011; Zhang et al., 2012). Maintaining SNARE and dynamin function is likely key to CSP’s neuroprotective role (Burgoyne and Morgan, 2011; Rozas et al., 2012; Sharma et al., 2012a; Sharma et al., 2011).

The *CLN4* causing dominant mutations L115R and L116Δ are clustered in the palmitoylated cysteine-string (CS) domain of CSPα, which mediates CSPα’s secretory trafficking to axon terminals, its SV association, and its dimerization (Arnold et al., 2004; Chamberlain and Burgoyne, 1998b; Greaves and Chamberlain, 2006; Greaves et al., 2008; Ohyama et al., 2007; Stowers and Isacoff, 2007; Swayne et al., 2003). Palmitoylation of the CS domain enables CSPα’s export from the ER and Golgi (Chamberlain and Burgoyne, 1998b; Greaves and Chamberlain, 2006; Greaves et al., 2008; Ohyama et al., 2007; Stowers and Isacoff, 2007). Palmitoylation must then be maintained for CSPα’s SVP trafficking and/or SV association, presumably to due to the short lifetime of palmitoylation (Fukata and Fukata, 2010). The latter has been indicated by much reduced synaptic levels CSP in loss of function mutants of the synaptic palmitoyl-transferase HIP14/DHHC17 (Ohyama et al., 2007; Stowers and Isacoff, 2007). Notably, there is a link between CSPα’s degree of lipidation and lysosomal dysfunction. In a lysosomal disease mouse model of Mucopolysaccharidosis type IIIA (MPS-IIIA) palmitoylation of CSPα was decreased and its proteasomal degradation was increased (Sambri et al., 2017). Since overexpression (OE) of CSPα in MPS-IIIA mice ameliorated their presynaptic defects, neurodegeneration, and prolonged survival, CSPα could be a critical factor for the progression of many lysosomal diseases (Sambri et al., 2017).

Post-mortem analysis of *CLN4* patient brains suggests that dominant *CLN4* mutations have two key pathological effects: to reduce monomeric levels of lipidated CSPα and promote the formation of high-molecular weight CSPα protein aggregates/oligomers that are ubiquitinated (Greaves et al., 2012; Henderson et al., 2016; Noskova et al., 2011). Similar effects of the mutations were seen in HEK293T, PC12 cells, and fibroblasts from *CLN4* carriers (Benitez and Sands, 2017; Greaves et al., 2012; Zhang and Chandra, 2014). *In vitro*, *CLN4* mutant CSPα aggregates form in a time-, concentration- and temperature-dependent manner (Zhang and Chandra, 2014). Palmitoylation of CSPα’s promotes aggregation (Greaves et al., 2012), although it is not essential (Zhang and Chandra, 2014). In addition, post-mortem brains of *CLN4* patients exhibit large scale changes in protein palmitoylation (Henderson et al., 2016). *CLN4* mutations have no adverse short-term effects on CSPα’s co-chaperone functions *in vitro*, including activation of Hsc70’s ATPase activity, or the binding to chaperone clients like SNAP25 and dynamin (Zhang and Chandra, 2014). However, during prolonged incubation, the ability of mutant CSPα to stimulate Hsc70’s ATPase declines considerably (Zhang and Chandra, 2014).

Several mechanisms of *CLN4* induced neurodegeneration have been suggested. As with other neurodegenerative diseases, progressive aggregation of CSPα may account for the dominant nature of *CLN4* mutations (Greaves et al., 2012; Henderson et al., 2016; Noskova et al., 2011; Zhang et al., 2012). In turn, this may cause a haplo-insufficiency that triggers neurodegeneration (Greaves et al., 2012; Noskova et al., 2011; Zhang and Chandra, 2014). An alternative gain of function mechanism involves mutant CSPα aggregation, which may have a dominant-negative effect and sequester WT CSPα or other proteins (Greaves et al., 2012; Henderson et al., 2016; Zhang et al., 2012). However, the link between *CLN4* mutations, lysosomal dysfunction and neurotoxicity remains unclear. Therefore, an animal model may provide critical insight into molecular and cellular mechanisms underlying *CLN4* disease.

Here, we generated to *Drosophila* models for *CLN4* by expressing by expressing either *CLN4* mutant hCSPα or dCSP in fly neurons human. The human model has the unique advantage of being able to separate effects of CLN4-mutant hCSPα from effects in WT dCSP. We show that both fly models replicate all key pathogenic biochemical properties of *CLN4* including decreased monomeric CSP levels and increased levels of high-molecular weight and ubiquitinated CSP oligomers. Further analysis revealed novel insights into mechanisms underlying *CLN4* pathology.

## Results

### Generation of a *Drosophila CLN4* model

We generated a *Drosophila* model of *CLN4* by expressing the disease-causing human proteins hCSPα-L115R, hCSPα-L116Δ (Fig. 1A; denoted as L115 and L116 from now on) and the corresponding WT hCSPα control from a common genomic phi31-attP insertion site under the transcriptional control of the yeast Gal4-UAS expression system (Brand and Perrimon, 1993). Unless otherwise indicated, we used the pan-neuronal elav-Gal4 driver C155 (Lin and Goodman, 1994) to express these proteins exclusively in otherwise WT neurons (*w^1118^*).

**Figure 1.**
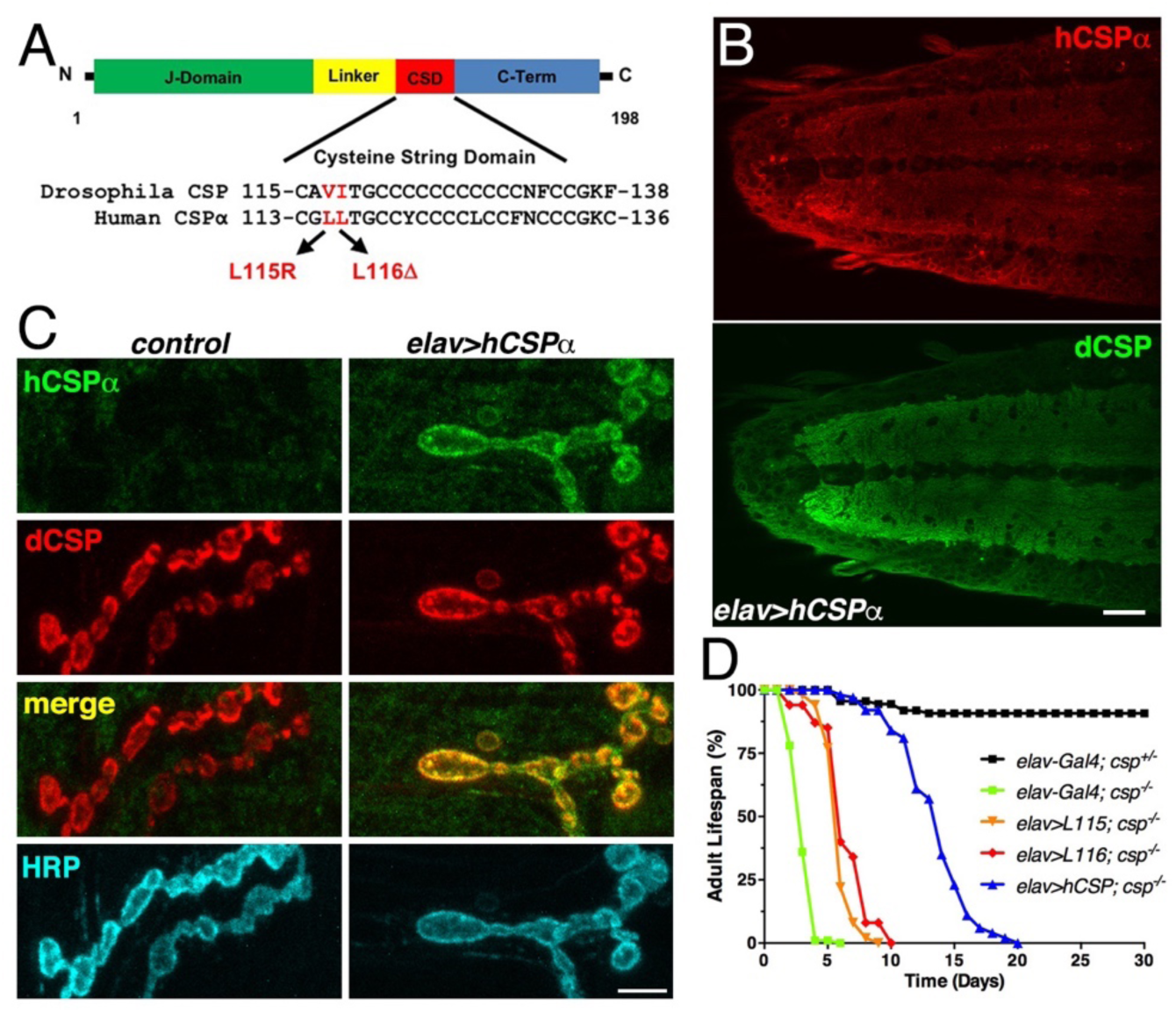
Generation of a *Drosophila CLN4* model. **A.** Structure of CSP and position of *CLN4* mutations in the CS domain of hCSPα and dCSP. CSP’s N-terminal J domain, linker domain and C-terminus are indicated. **B.** Larval VNC of animals expressing WT hCSPα in neurons from a single transgene with an elav driver immunostained for hCSPα and endogenous dCSP. Scale bar, 20 μm. **C.** Larval NMJs of control and animals expressing WT hCSPα immunostained for hCSPα, dCSP, and HRP marking the presynaptic plasma membrane. Scale bar, 5 μm. **D**. Adult lifespan of control (*dcsp^X1/+^*, black), *dcsp^X1/R1^* deletion mutants (green) and mutants expressing WT hCSPα (blue), hCSP-L115, (orange), or -L116 (red) with an elav driver.

To evaluate whether WT hCSPα is at least partially functional in fly neurons, we first determined whether neuronally expressed hCSPα is properly palmitoylated and targeted to SVs. This analysis was aided by the availability of species-specific antibodies that discriminate between hCSPα and dCSP (Figs. 1B-C, 2A; S1C, S2A). Neuronally expressed hCSPα was efficiently trafficked to axon terminals and co-localized with endogenous dCSP in the neuropil of the larval ventral nerve cord (VNC; Fig. 1B) and larval neuromuscular junctions (NMJs; Fig. 1C). Consistent with the normal trafficking of hCSPα, the majority of neuronally expressed hCSPα was fully palmitoylated (Fig. 2A), which was confirmed by treating larval protein extracts with 0.5 M hydroxylamine (Fig. S1A).

**Figure 2.**
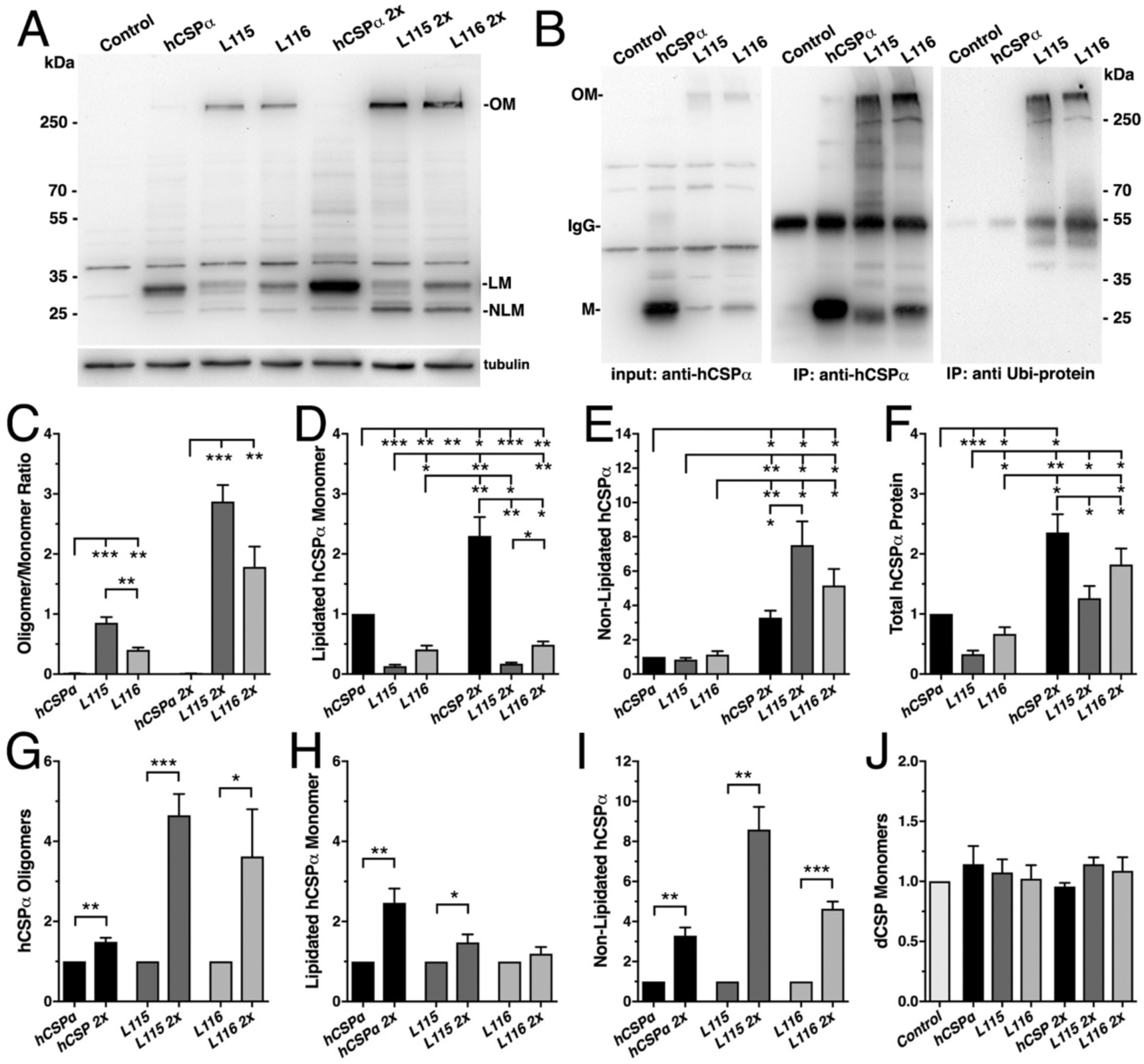
*CLN4* mutations cause dose-dependent oligomerization hCSPα in neurons. WT and mutant hCSPα (L115/L116) were expressed in larval neurons of *white^1118^* animals (control) with an elav driver from 1 or 2 transgenes (2x). **A.** Western blot of larval brain protein extracts probed for hCSPα. Signals for SDS-resistant hCSPα oligomers (OM), lipidated monomeric hCSP (LM), and non-lipidated hCSP (NLM) are indicated. β-tubulin was used as loading control. **B.** Western blots probed for hCSPα or lysine-linked-ubiquitin of hCSPα-immunoprecipitated extracts from adult heads of indicated genotypes. Signals for IgG heavy chain are indicated. **C.** Average oligomer/monomer ratios (N = 5). **D-F.** Levels of lipidated (D), non-lipidated (E), and total hCSPα (F) normalized to WT hCSPα (N = 6). **G-I.** Dosage-dependent increase of hCSPα oligomer (G), lipidated (H), and non-lipidated monomer levels (I). Signals were normalized to loading control and plotted as n-fold change from 1-copy expression of WT hCSPα (N = 6). **J.** Levels of monomeric dCSP shown as n-fold change from control (N = 3). Graphs display mean ± SEM. Statistical analysis used one-way ANOVA (C-F, J) or two-tailed unpaired *t* test (G-I); *, P < 0.05; **, P < 0.01; ***, P < 0.001.

Next, we tested whether hCSPα can functionally replace endogenous dCSP and restore the premature lethality of flies lacking dCSP (Zinsmaier et al., 1994). Flies and mice lacking CSP due to a gene deletion exhibit progressive neurodegeneration, paralysis and premature death (Fernandez-Chacon et al., 2004; Zinsmaier et al., 1994). Pan-neuronal elav-driven expression of normal hCSPα from one transgenic copy significantly restored adult lifespan of homozygous *dcsp* deletion mutants from ∼4-5 days to 15-20 days (LD50 p<0.001; Fig. 1D). Even though this rescue was only partial, it confirms that hCSPα is functional in flies, which made a *CLN4 Drosophila* model using human proteins tenable.

Notably, neuronal expression of *CLN4* mutant hCSPα in a *dcsp* null background was able to partially rescue the lifespan deficit (p<0.001, LD50), although not nearly to the extent of wild type hCSPα (Fig. 1D). This indicates that *CLN4* mutant hCSPα proteins are at least partially functional.

### *CLN4* mutations cause formation of SDS-resistant and ubiquitinated hCSPα protein oligomers in neurons

In humans, *CLN4* mutations reduce levels of lipidated monomeric hCSPα and drive the formation of high-molecular weight, SDS-resistant hCSPα protein oligomers that are ubiquitinated (Benitez et al., 2015; Greaves et al., 2012; Henderson et al., 2016; Noskova et al., 2011). Both features were also present when mutant hCSP-L115 or -L116 were expressed in non-neuronal and neuronal mammalian cell cultures with L115 exhibiting stronger effects than L116 (Diez-Ardanuy et al., 2017; Greaves et al., 2012; Henderson et al., 2016; Zhang and Chandra, 2014).

To test whether expression of *CLN4* mutant hCSPα in *Drosophila* neurons recapitulates the pathological features of post-mortem human brains, we analyzed the properties of *CLN4* mutant hCSPα by immunoblotting, using protein extracts from larval VNCs. When expressed from a single transgene with an elav-Gal4 driver, lipidated monomeric protein levels of WT hCSPα were about 0.8-1.5 times of endogenous dCSP levels (data not shown). In comparison, levels of lipidated monomeric hCSP-L115 and -L116 were significantly reduced to 13% and 40% of WT hCSPα levels, respectively (p<0.01, Fig. 2A, D). Levels of lipidated hCSP-L115 monomers were significantly lower than hCSP-L116 (p<0.05, Fig. 2D). Levels of non-lipidated monomeric hCSP-L115 and -L116 were comparable to WT hCSPα levels (p>0.2; Fig. 2A, E).

Both *CLN4* mutations induced the formation of SDS-resistant, high-molecular weight hCSPα oligomers in *Drosophila* neurons (p<0.01; Fig. 2A, C). In contrast, WT hCSPα oligomers were barely detectable (Fig. 2A, C). The mutation L115 triggered oligomerization to a significantly larger degree than L116 (p<0.01; Fig. 2C), which is consistent with the lower levels of hCSP-L115 monomers (Fig. 2D). Increasing levels of DTT in the buffer reducing disulfide bonds had little to no effect on the levels and size of hCSP-L115 and -L116 oligomers, which varied widely from ∼250 kDa to more than 500 kDa (not shown).

To determine whether *CLN4* mutant hCSPα oligomers are ubiquitinated, we immunoprecipitated hCSPα from larval brain cell lysates and probed western blots with an antibody that specifically detects K29-, K48-, and K63-linked mono- and poly-ubiquitinated proteins but not free ubiquitin monomers (Fujimuro et al., 1994). hCSPα antibodies immunoprecipitated both hCSPα monomers and oligomers (Fig. 2B) but not endogenous dCSP (not shown). The anti-ubiquitinated protein antibody only recognized a strong ∼250 kD protein band in precipitates from L115 and L116 mutant brains, which correlated in size with high-molecular weight hCSPα oligomers (Fig. 2B). No ubiquitin-positive signals corresponding to *CLN4* mutant hCSPα monomers, WT hCSPα monomers or oligomers were detected (Fig. 2B). Taken together, these data suggest that expression of *CLN4* mutant hCSPα in fly brains reproduces the critical biochemical pathological features of post-mortem human *CLN4* brains (Greaves et al., 2012; Henderson et al., 2016; Noskova et al., 2011).

### Oligomerization of *CLN4*-mutant hCSPα is dose-dependent

Oligomerization of purified *CLN4* mutant hCSPα proteins is dose-dependent *in vitro* (Zhang and Chandra, 2014). To test whether this is also the case *in vivo*, we doubled mRNA levels of WT and *CLN4*-mutant hCSPα by expressing two copies of the respective transgenes with a single elav-Gal4 driver. In comparison to the expression from one transgene, levels of lipidated and non-lipidated monomeric WT hCSPα increased ∼2.4-fold and ∼3.3-fold, respectively (p<0.01, Fig. 2A, D-E, H-I). Yet, WT hCSP oligomer levels increased only 1.5-fold (p<0.01; Fig. 2G) and remained low such that the oligomer/monomer ratio was not altered (p=0.9; Fig. 2C).

In comparison to single copy gene expression, doubling gene dosage increased levels of lipidated hCSP-L115 monomers only modestly by 1.5-fold (p<0.05) while hCSP-L116 levels were not significantly affected (p=0.3; Fig. 2H). However, levels of hCSP-L115 and -L116 oligomers increased 4.6- and 3.6-fold, respectively (p<0.05; Fig. 2G). Due to the disproportional increase of *CLN4* mutant monomers and oligomers, the oligomer/monomer ratio increased from 0.9 to 2.8 for hCSP-L115 and from 0.4 to 1.8 for hCSP-L116 (Fig. 2C). Hence, oligomerization of hCSP-L115 and -L116 is dose-dependent.

Doubling mRNA expression disproportionally increased levels of non-lipidated hCSP-L115 and - L116 monomers 8.6- and 4.6-fold, respectively (p<0.01, Fig. 2I). The increase of non-lipidated mutant monomers is unlikely due to a rate limiting effect on mechanisms mediating palmitoylation because levels of non-lipidated WT monomers increased much less than mutant monomers, even though overall levels of lipidated WT monomers were much higher (Fig. 2E).

Doubling gene dosage of hCSP-L115 and -L116 expression increased total overall protein levels 3.0 and 3.9-fold, respectively (p<0.01; Fig. S1D). It also preserved the relative difference in overall protein levels between WT and *CLN4* mutants (Fig. 2F). Since lipidated mutant monomer levels remained unaltered (p>0.1; Fig. 2D), the increase in overall protein levels of *CLN4* mutant hCSPα is essentially due to increased levels of oligomers and non-lipidated monomers (p<0.05; Fig. 2C, E).

Levels of endogenous dCSP monomers were not affected by low or high expression of *CLN4* mutant hCSPα (Fig. 2J). Notably, high-molecular weight oligomers of endogenous WT dCSP were not detectable after expression of *CLN4* mutant hCSPα (Fig. S1C), which contrasts a previous study detecting small amounts of overexpressed GFP-tagged WT hCSPα in *CLN4* mutant hCSPα oligomers (Greaves et al., 2012).

### Oligomerization of neuronally expressed *CLN4*-mutant hCSPα precedes lethality

Elav-driven neuronal expression of WT or *CLN4*-mutant hCSPα from a single transgene had no effect on viability during development (Fig. 3A) and adult lifespan (not shown). However, doubling gene expression levels severely reduced viability of hCSP-L115 and -L116 animals in comparison to WT hCSPα expression (p<0.001; Fig. 3A). There was no significant difference between L115 and L116 mutant animals (p=0.6; Fig. 3A). The sharp drop in viability induced by 2-copy expression of hCSP-L115 and -L116 indicates a tight threshold of neurotoxicity, which might be linked to the more than 3.6-fold increase of *CLN4* mutant CSPα oligomers and/or the more than 4.6-fold increase of non-lipidated monomers (Fig. 2G, I).

**Figure 3.**
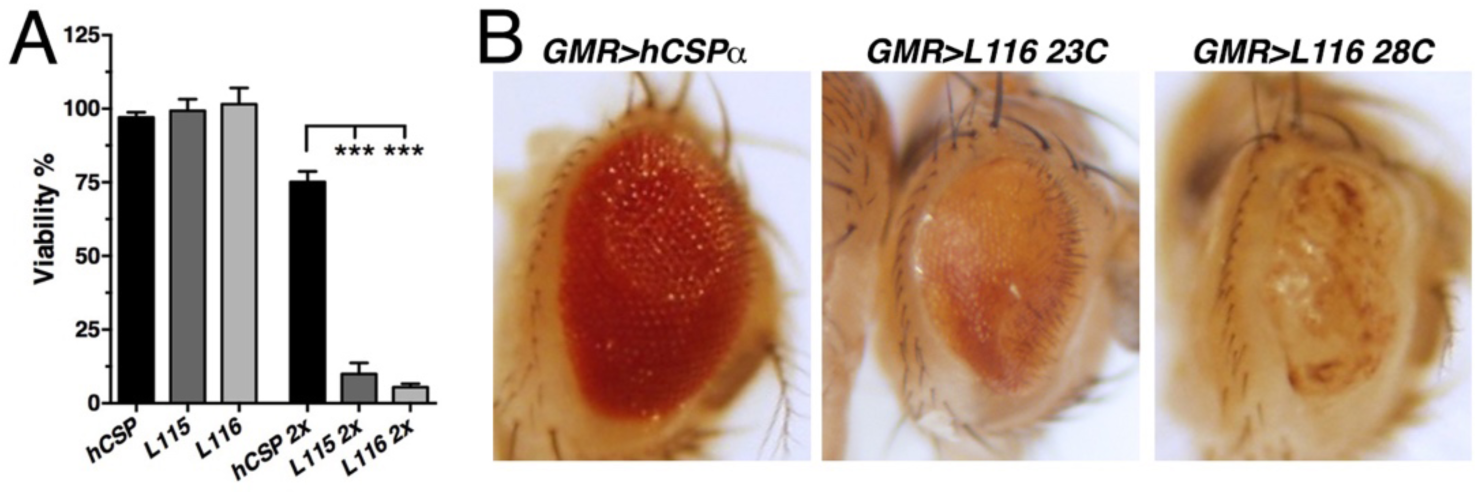
*CLN4* mutations cause dose-dependent lethality and eye degeneration. **A.** Viability of animals expressing WT, L115, or L116 mutant hCSPα pan-neuronally from 1 or 2 transgenes (2x) with an elav driver (mean ± SEM; N ≥ 3, n ≥ 740; ***, P < 0.001, one-way ANOVA). **B.** Images of adult fly eyes expressing WT hCSPα or hCSPα-L116 with a GMR-Gal4 driver at 23ºC and 28ºC.

Expression of *CLN4*-mutant hCSPα with the pan-neuronal nSyb-Gal4 driver also caused a dose-dependent lethality, which was similar to that of the elav-Gal4 driver (data not shown). In comparison to the elav-driven expression, *CLN4*-mutant adult escapers were more sluggish and hardly moved. They were also unable to properly inflate their wings (Fig. S1B) and died prematurely within a few days.

To further examine potential degenerative effects in adults, we exclusively expressed hCSP-L116 in the eye by using the eye-specific GMR-Gal4 driver (Hay et al., 1997). High-level expression of WT hCSPα from a line with multiple transgene insertions (see methods) had essentially no effect at 23°C or 28°C (Fig. 3B). In contrast, expression hCSP-L116 at 23°C severely impaired the size, integrity and pigmentation of the eye (Fig. 3B). Raising flies at 28°C to increase Gal4 activity and thereby gene expression enhanced the severity of the degenerative phenotype of L116-mutant eyes (Fig. 3B). This enhancement is consistent with a dose-dependent increase in hCSP-L116 oligomerization (Fig. 2C, G) but the higher temperature may also increase misfolding and/or oligomerization of mutant hCSPα.

### *CLN4* mutations impair hCSPα’s synaptic localization

Previously, it has been suggested that *CLN4* mutations may disrupt anterograde trafficking of CSPα, which could be due either to a direct effect on the palmitoylation of hCSP monomers or oligomer formation (Benitez et al., 2011; Greaves et al., 2012; Noskova et al., 2011). However, whether *CLN4* mutations indeed affect the synaptic localization of CSPα remained unclear. To assess this, we triple-immunostained 3^rd^ instar larval NMJs with antibodies against hCSPα, endogenous dCSP marking SVs, and HRP marking the neuronal plasma membrane. In comparison to WT hCSPα, levels of mutant hCSP-L115 and -L116 were significantly reduced at synaptic boutons (p<0.05; Fig. 4A, H). Expression of *CLN4* mutant or WT hCSPα had no effect on synaptic levels of endogenous WT dCSP at NMJs (Fig. 4A, I). Despite the reduced protein levels, mutant hCSPα still partially rescued the lifespan deficit of *dcsp* deletion mutants (p<0.001, LD50), although not nearly to the extent of WT hCSPα (Fig. 1D). This indicates that *CLN4* mutant hCSPα proteins are at least partially functional.

**Figure 4.**
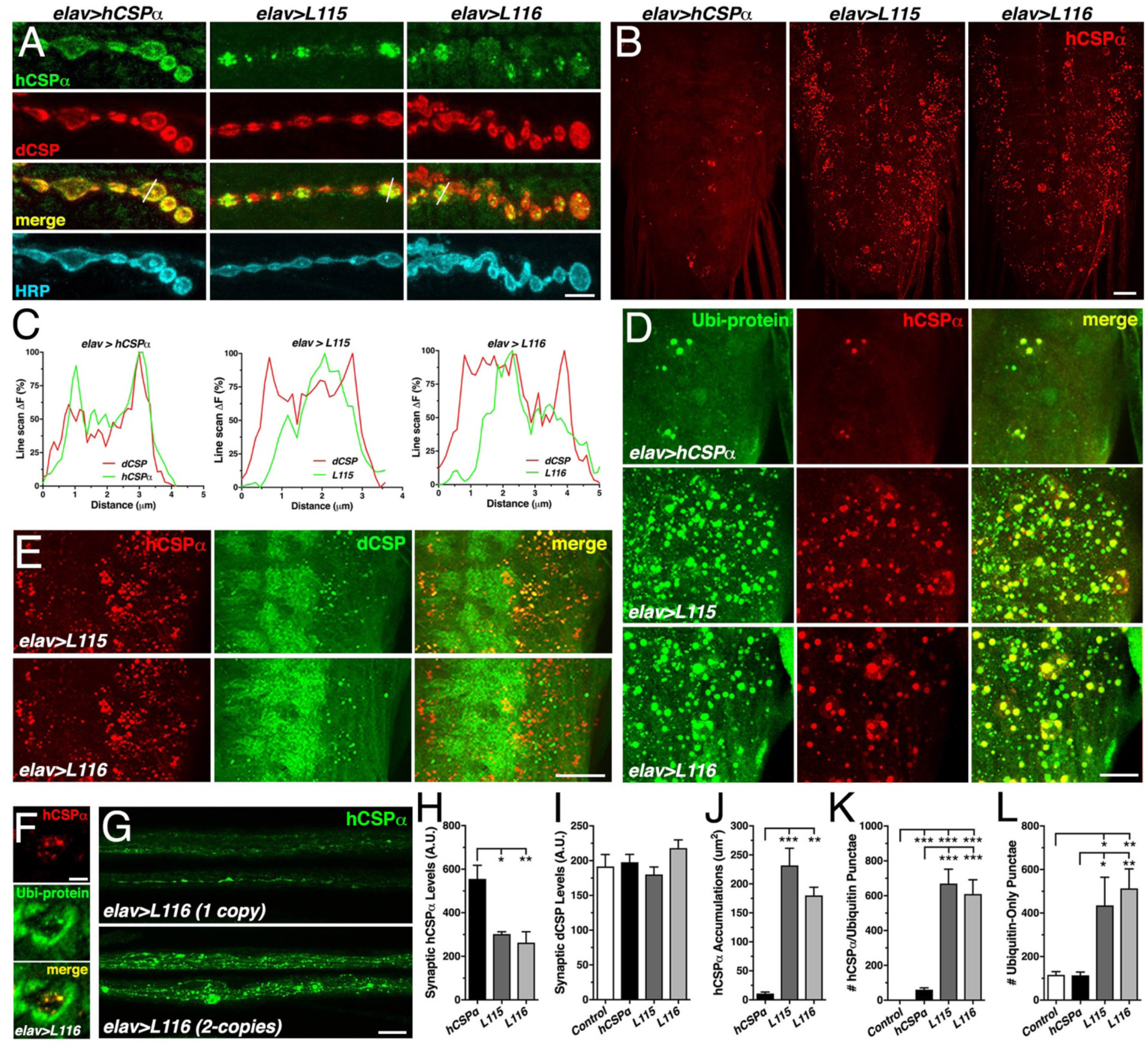
*CLN4* mutations reduce synaptic hCSPα levels and cause abnormal accumulations with endogenous dCSP and ubiquitinated proteins in axons and somata. WT, L115 or L116 mutant hCSPα were expressed in larval neurons with an elav-Gal4 driver from one (D-E, J-L) or two (A, H-I) transgenes. Genotypes are indicated. **A.** Larval NMJs immunostained for hCSPα, endogenous dCSP, and the neuronal membrane marker HRP. Dashed lines denote line scans shown in C. **B.** Larval VNCs stained for hCSPα. **C.** Plots of hCSPα and dCSP fluorescence from single line scans through synaptic boutons (dashed lines in A). **D.** Larval VNC segments stained for hCSPα (red) and lysine-linked-ubiquitin visualizing ubiquitinated proteins (Ubi-protein). **E.** Larval brain segments stained for hCSPα and dCSP. **F**. Synaptic bouton of larval NMJ stained for hCSPα and Ubi-proteins. **G.** Proximal larval segmental nerves stained for hCSPα. **H-I.** Average levels of hCSPα (H) and dCSP (I) at synaptic boutons of larval NMJs (N > 4). **J.** Cumulative area of abnormal hCSPα accumulations in larval brains (N ≥ 3). **K-L.** Average number of accumulations immunopositive for both hCSPα and ubiquitin (K), or only positive for ubiquitin (L) but not hCSPα (N ≥ 4). Scale bars: 5 μm (A), 20 μm (B, G), 15 μm (D), 10 μm (E), 5 μm (F). Graphs display mean ± SEM. Statistical analysis used one-way ANOVA (H-L); *, P < 0.05; **, P < 0.01; ***, P < 0.001.

WT hCSPα co-localized with endogenous dCSP uniformly in the periphery of synaptic boutons (Fig. 4A, C). In contrast, hCSP-L115 and -L116 were enriched in brightly stained clusters that were more distant from the presynaptic membrane (Fig. 4A, C). In comparison to control, *CLN4* mutant hCSPα also accumulated abnormally in axons of segmental nerves (Fig. 4G) and the larval brain (p<0.01; Fig. 4B, I, S2A). The size of *CLN4* mutant hCSPα accumulations in the larval brain was heterogeneous, ranging from ∼500 nm in diameter to ∼3 μm in neuronal somata. Occasionally, extreme accumulations of mutant hCSPα were also observed at synaptic boutons of larval NMJs (Fig. S2C). The occurrence of abnormal hCSP-L115 and -L116 accumulations was also dose-dependent (not shown), which 0was most pronounced in axons of segmental nerves (Fig. 4G). Hence, *CLN4* mutations not only reduce synaptic levels of hCSPα but also cause a severe mislocalization.

### Abnormal accumulations of *CLN4* mutant hCSPα contain WT dCSP and ubiquitinated proteins

A fraction of the abnormal accumulations of hCSP-L115 and -L116 contained significant amounts of endogenous WT dCSP in axons (not shown) and somata (Fig. 4E, S2A). Since WT dCSP was not detected together with high-molecular weight oligomers of *CLN4* mutant hCSPα on Western blots, this indicated that *CLN4* mutant hCSPα and WT dCSP may co-accumulate on intracellular membranes.

Theoretically, *CLN4* mutant hCSPα could accumulate on ER, Golgi, or abnormal SVP membranes during secretory trafficking. Alternatively, mutant hCSPα could accumulate on endosomal membranes that are targeted for degradation. To test whether mutant hCSPα accumulations contain ubiquitinated proteins indicative of prelysosomal membranes, we used antibodies that exclusively detect ubiquitinated proteins but not monomeric ubiquitin (Fujimuro et al., 1994). Indeed, essentially all *CLN4* mutant hCSPα accumulations were immunopositive for ubiquitinated proteins in somata and synaptic boutons of NMJs (Fig. 4D, F, S2B). In comparison to larvae expressing WT hCSPα, the amount of hCSPα accumulations containing ubiquitinated proteins was significantly increased in *CLN4* mutants (p<0.001; Fig. 4K). Hence, the abnormal accumulations of *CLN4* mutant hCSPα are either enriched in ubiquitinated hCSPα oligomers and/or contain other ubiquitinated proteins that are potentially destined for protein degradation.

### *CLN4* mutations impair protein homeostasis

The co-immunostainings against hCSPα and ubiquitinated proteins also revealed a substantial increase in the levels of ubiquitinated proteins that were not associated with accumulations of *CLN4* mutant hCSPα (Fig. 4D; S2B). Brains of control (*w^1118^*) and larvae expressing WT hCSPα contained a similar amount of ubiquitinated protein foci that were negative for hCSPα (Fig. 4L), indicating that expression of WT hCSPα exerts little to no effect on protein ubiquitination. In comparison, the amount of ubiquitinated protein foci that were not associated with mutant hCSPα was significantly increased in brains expressing hCSPα-L115 or -L116 (p<0.05; Fig. 4L). Hence, *CLN4* mutant hCSPα may directly or indirectly affect a step of protein homeostasis that that leads to excessive protein ubiquitination of unrelated proteins.

### *CLN4* mutations in fly and human CSP have similar effects

To validate the observed phenotypes of *CLN4* mutant hCSPα and exclude that they are not an artefact of expressing mutant human proteins in fly neurons, we expressed *CLN4* mutant dCSP2 (dCSP2) containing the mutations V117R and I118Δ, which are analogous to the mutations L115R and L116Δ in hCSPα (Fig. 1A). All fly transgenes were expressed pan-neuronally with an elav-Gal4 driver from the same transgenic insertion site as the human transgenes. Expression levels of monomeric lipidated dCSP2-V117 and -I118 were reduced in comparison to WT (Fig. S3A). Similar to its human analog L115 (Fig. 2A), V117 reduced levels of monomeric dCSP more severely (Fig. S3A). In addition, both V117 and I118 triggered the formation of SDS-resistant, high-molecular weight oligomers (Fig. S3A), which were absent for WT dCSP2 expression. Expressing dCSP2-V117 and -I118 in *dcsp* deletion mutants had similar effects (Fig. S3A). Hence, the pathological features of the dominant *CLN4* mutations reducing levels of lipidated CSP monomers and triggering protein oligomerization are conserved between fly and human CSP.

To determine the subcellular localization of *CLN4* mutant dCSP in fly neurons, we expressed normal and *CLN4* mutant proteins from a single transgene in neurons of homozygous *dcsp* deletion null mutants. Like for mutant hCSPα, levels of dCSP-V117 and -I118 were reduced at synaptic boutons of larval NMJs and the neuropil of the larval VNC (Fig. S3B-C). Mutant dCSP was also mislocalized and accumulated abnormally in axons and neuronal somata of the larval brains (Fig. S3B).

### *CLN4* mutant hCSPα accumulates on prelysosomal endosomes

To further define the nature of the co-accumulations of *CLN4* mutant hCSPα with ubiquitinated proteins, we tested a number of organelle markers for a potential colocalization. Mutant hCSPα accumulations did not colocalize with the endoplasmic reticulum (ER), cis-, or trans-Golgi complexes (Fig. 5A), which excluded a major defect in ER or Golgi trafficking.

**Figure 5.**
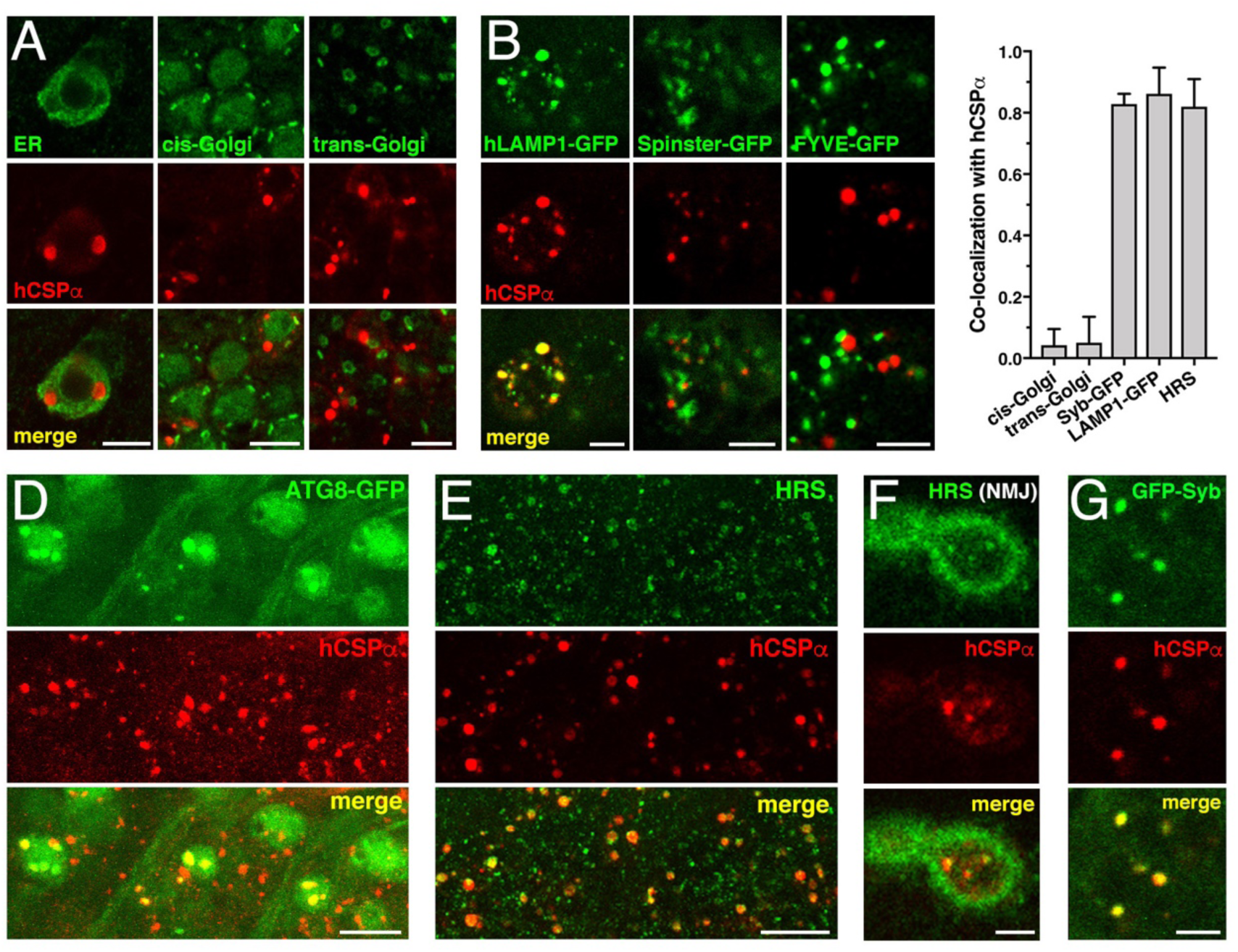
*CLN4* mutant hCSPα accumulates on LAMP1- and HRS-positive endosomes. hCSP-L116 was expressed in larval neurons with an elav-Gal4 driver from 1 transgene. As indicated, respective reporter transgenes were co-expressed. **A.** Neurons of larval VNCs co-immunostained for hCSPα and the ER marker GFP-KDEL, the cis-Golgi marker GMAP, or the trans-Golgi marker Golgin 245. **B.** Neurons co-immunostained for hCSPα (red) and co-expressed hLAMP1-GFP, Spinster-GFP or the PI_3_P marker FVVE-GFP. **C.** Fraction of organelle markers colocalizing with hCSPα accumulations (mean ± SEM; n ≥ 65, N ≥ 4). **D-E.** Segments of larval brains costained for hCSPα and ATG8/LC3-GFP (D) or HRS (E). **F.** Synaptic boutons at larval NMJs co-stained for hCSPα and HRS. **G.** Neuron co-immunostained for hCSPα and coexpressed GFP-nSyb. Scale bars: 10 μm (D-E), 5 μm (A-B, F-G).

A large fraction of mutant hCSPα co-accumulated with hLAMP1-GFP (Fig. 5B-C), which labels a heterogeneous population of organelles ranging from pre-degradative endosomal species to degradative lysosome (Cheng et al., 2018; Saftig and Klumperman, 2009; Yap et al., 2018). A similar large fraction of mutant hCSPα also co-accumulated with the hepatocyte growth factor regulated tyrosine kinase substrate (HRS, Fig. 5C, E), which is a critical component of “endosomal sorting complexes required for transport” (ESCRT) mediating the transition from early to late endosomes (Raiborg and Stenmark, 2009). Only a small fraction co-localized with coexpressed Rab5-GFP (not shown) or the autophagosomal marker ATG8/LC3-GFP (Fig. 5D). No co-localization was observed with lysosomal Spinster-GFP (Fig. 5B; (Rong et al., 2011; Sweeney and Davis, 2002) or the late endosomal protein Rab7 (not shown; (Guerra and Bucci, 2016).

Notably, endosomes accumulating mutant hCSPα did not co-localize with the co-expressed phosphatidylinositol 3-phosphate (PI_3_P) sensor FYVE-GFP (Fig. 5B), even though HRS requires PI_3_P for membrane association (Mayers et al., 2013; Raiborg et al., 2001b). Hence, this suggests that all PI_3_P binding sites may be occupied on the abnormal endosomes accumulating mutant hCSPα. Alternatively, PI_3_P may be absent from these endosomes, which would imply an abnormal retention of HRS. Taken together, these data suggest that *CLN4* mutant hCSPα accumulates on prelysosomal endosomes that are inefficiently processed for lysosomal fusion.

Consistent with the colocalization of mutant hCSPα with ubiquitinated proteins at axon terminals (Fig. 4F), HRS also colocalized with mutant hCSPα at synaptic boutons of NMJs (Figs. 5F). This raised the possibility that hCSPα-positive endosomes may originate from synaptic terminals. Consistently, hCSPα-positive accumulations co-localized with SV-associated Synaptobrevin-GFP (Syb-GFP; Fig. 5C, G) while a small subset co-localized with the synaptic plasma membrane protein Syntaxin 1A (not shown). The CSPα chaperone clients Dynamin and SNAP-25 (Sharma et al., 2011; Zhang et al., 2012) did not co-localize with hCSPα-positive endosomes (not shown).

### *CLN4* mutations cause various ultrastructural endo-membrane abnormalities in axons and neuronal somata

Since *CLN4* mutant hCSPα abnormally accumulates on prelysosomal endosomes, we used electron microscopy to detect potential ultrastructural defects in membrane trafficking. Expression of hCSPα-L115 and -L116 induced highly abnormal membrane structures in neuronal somata, the neuropil of the larval VNC, and axons of segmental nerves (Fig. 6). The most prominent and frequent abnormal structures were multilamellar “membrane whirls” of various shapes and density that contain highly electron-dense membranes (arrowheads, Fig. 6A-B, C-D, F-G). Like endosomal hCSPα-positive accumulations detected by confocal microscopy, membrane whirls were most frequently observed in the neuropil of the VNC and axons of segmental nerves (Fig. 6C-D). To a lesser degree, they were present in the cytoplasm of neuronal somata (Fig. 6A-B, F-G) and occasionally the nucleoplasm (not shown). Sporadically, whirls were found on opposite sides of plasma membranes of neighboring cells (arrowhead, Fig. 6B).

**Figure 6.**
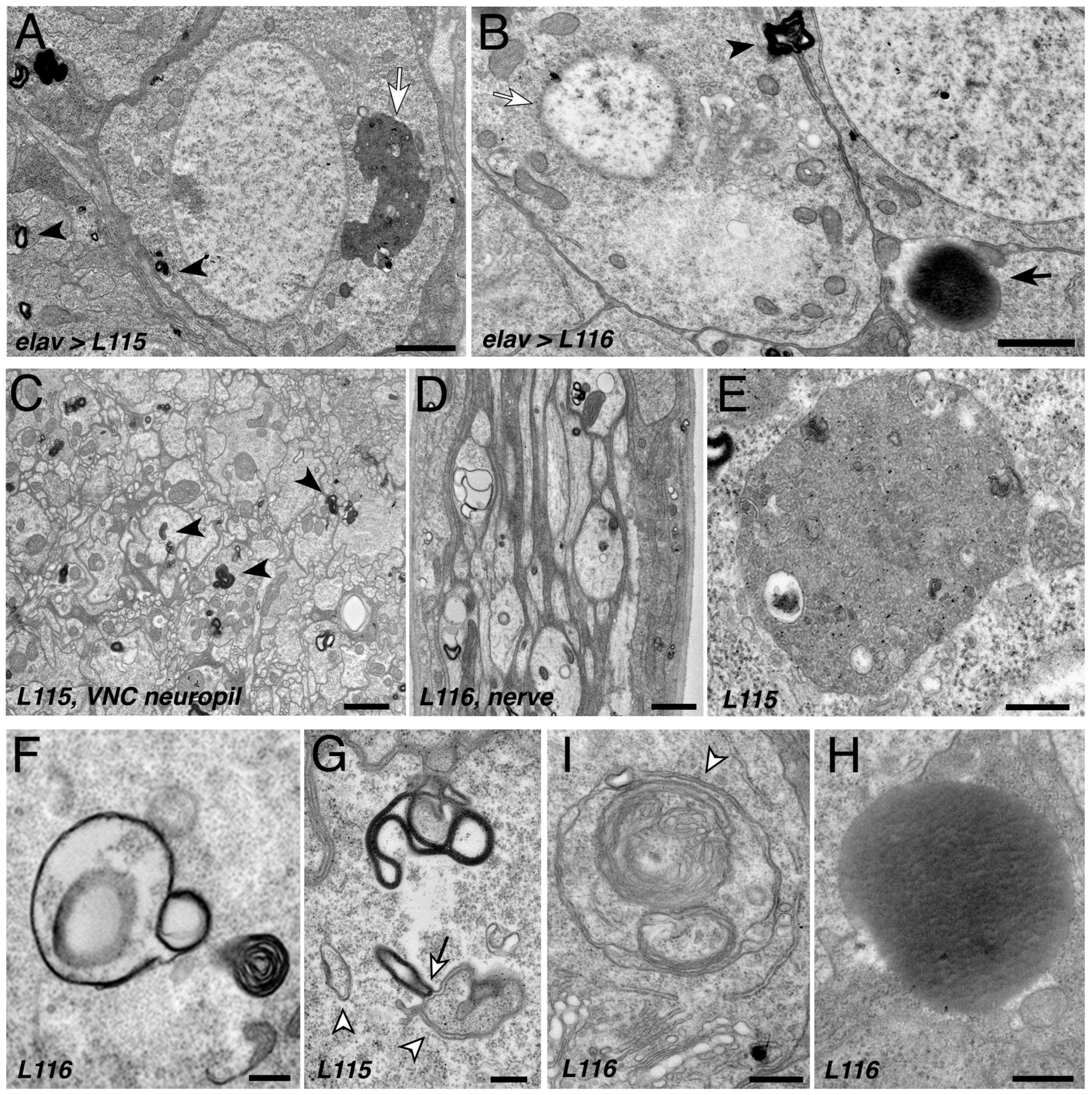
*CLN4* mutations cause abnormal endomembrane structures and EM-dense accumulations. TEM micrographs of ultrathin (70 nm) sections from larval VNCs expressing hCSP-L115 or L116 with an elav driver from 2 transgenic copies. Genotypes are indicated. **A-B.** Neuronal somata containing electron-dense membrane whirls (black arrowheads), large electron-dense extracellular deposits (black arrow, B), and occasionally “residual lysosomes” (white arrow, A), bloated Golgi Apparati (B), and degenerating nuclear membranes (white arrow, B). **C.** Neuropil of larval VNC containing membrane whirls in neuronal processes (black arrowheads). **D.** Sagittal section of larval segmental nerve containing membrane whirls and abnormal autophagosome-like structures in axons of sensory and motor neurons. **E-H**. High magnification images showing residual lysosome with a diverse variety of intraluminal vesicles (E), various forms of EM-dense membrane whirls (F-G) and autophagosome-like structures (white arrowheads, G-I) that may interact with EM-dense structures (white arrow, G), and an electron-dense extracellular deposit (H). Scale bars: 1 μm (A-C), 200 nm (D-E).

In addition to electron-dense membrane whirls, a number of secondary (residual) lysosomes (Fig. 6A, E) and abnormal autophagosome- and/or amphisome-like structures were present (grey arrowhead, Fig. 6F-I). Interestingly, EM-dense membranes forming whirls may interact with autophagosome (arrow, Fig. 6G). Notably, somewhat similar “membrane whirl” and autophagosome-like structures were observed in various ESCRT loss of function mutants including TSG101, Snf7/CHMP4B, and VPS4 and CHMP2B (Doyotte et al., 2005; Lee et al., 2007; Razi and Futter, 2006).

Next to abnormal membranes structures, homogenous electron-dense accumulations of unknown nature were frequently present in cellular regions of *CLN4* mutant larval brains (arrow, Fig. 6B, H). A limiting membrane was not detectable for these EM-dense accumulations, which are reminiscent but not identical of granular osmophilic deposits (GRODs) seen in post-mortem human tissue of *CLN4* patients (Anderson et al., 2013; Burneo et al., 2003; Noskova et al., 2011; Virmani et al., 2005). Since these structures were absent from the VNC neuropil and segmental nerves, they are unlikely a correlate for the abnormal endosomal accumulations of mutant hCSPα in axons.

Finally, a number of neurons that contained abnormal membrane structures also exhibited a severe fragmentation of the nuclear envelope and bloated Golgi cisternae (Fig. 6B), which together likely indicate a late stage of neuronal dysfunction and neurodegeneration (Nixon, 2006).

### Loss of ESCRT function causes endosomal accumulations of endogenous dCSP but no oligomerization

To verify that WT CSP is normally trafficked through the endo-lysosomal pathway, we impaired ESCRT function by an RNAi-mediated knock down (KD) of tumor suppressor gene 101 (TSG101), which is part of the ESCRT-I complex acting downstream of HRS and required for late endosome formation (Doyotte et al., 2005; Razi and Futter, 2006). Neuronal KD of TSG101 caused a mislocalization of endogenous dCSP, which accumulated on HRS-positive endosomes in the neuropil and neuronal somata of the larval VNC (Fig. S4A). Abnormal dCSP accumulations were also present in central regions of synaptic boutons of larval NMJs (Fig. S4B). Hence, WT CSP is likely being degraded though the endo-lysosomal pathway. This conclusion is consistent with the low amounts of lysosome-associated CSPα in human fibroblasts, N2A cells, mouse neurons, and lysosome-enriched fractions (Benitez and Sands, 2017; Chapel et al., 2013; Schroder et al., 2007; Tharkeshwar et al., 2017).

Notably, KD of TSG101 essentially mimicked the abnormal endosomal accumulation of *CLN4* mutant hCSPα-L115 and -L116 with one important exception. In contrast to hCSPα-L115 and -L116 expression, KD of TSG101 did not cause the formation of high-molecular weight dCSP oligomers (Fig. S4C), indicating that CSP oligomer formation is independent of impaired ESCRT function. In contrast to a recent report showing that impaired lysosomal activity in mouse neurons decreases CSPα’s palmitoylation (Sambri et al., 2017), impaired prelysosomal ESCRT function had no effect on the palmitoylation state of dCSP.

### The *CLN4* alleles hCSP-L115 and -L116 are hypermorphic gain of function mutations

Several hypotheses regarding the genetic nature of the dominant *CLN4* mutations have been suggested. Mutant hCSPα monomers and/or oligomers could acts as dominant-negatives sequestering normal CSP causing a haplo-insufficiency that triggers neurodegeneration (Greaves et al., 2012; Noskova et al., 2011; Zhang and Chandra, 2014). Alternatively, *CLN4* mutant alleles may induce toxicity via a gain of function (GOF) mechanism (Greaves et al., 2012; Henderson et al., 2016; Zhang et al., 2012). Such a GOF mutation could increase a normal activity of the protein (hypermorphic mutation) or introduce a new activity (neomorphic mutation). In theory, either type of mutation could trigger neurotoxicity.

To genetically address the genetic nature of *CLN4* mutations, we examined to what degree alterations of WT dCSP or hCSPα levels may affect L115 and L116 phenotypes. The rationale behind these genetic experiments is simple: lowering WT levels should enhance dominant-negative phenotypes, reduce hypermorphic phenotypes that are due to an increased normal activity, or have no effect on neomorphic phenotypes that are due a new protein activity (Muller, 1932; Wilkie, 1994).

Reducing endogenous WT dCSP levels by expressing two transgenic copies of *CLN4* mutant hCSP in heterozygous *dcsp* deletion mutants significantly suppressed the lethality induced by *CLN4* mutations (p<0.05, Fig. 7A). Conversely, increasing levels of WT hCSPα enhanced lethality (Fig. 7B-C). While expression of either WT or *CLN4* mutant hCSPα from a single transgene had no effect, co-expression of WT hCSPα with either one copy of hCSP-L115 or -L116 significantly reduced viability to ∼54% and 48%, respectively (p<0.001, Fig. 7B-C). The lethality induced by co-expression of WT and *CLN4* mutant hCSPα was significantly higher than the lethality induced by 2 copy expression of WT hCSPα (p<0.05, Fig. 7B-C). Hence, the modulating effects of altered WT dCSP or hCSPα levels on L115- and L116-induced lethality essentially exclude the possibility that either mutation acts as dominant-negative. Instead, these effects are consistent with the genetic characteristics of a hypermorphic GOF mutation (Muller, 1932; Wilkie, 1994).

**Figure 7.**
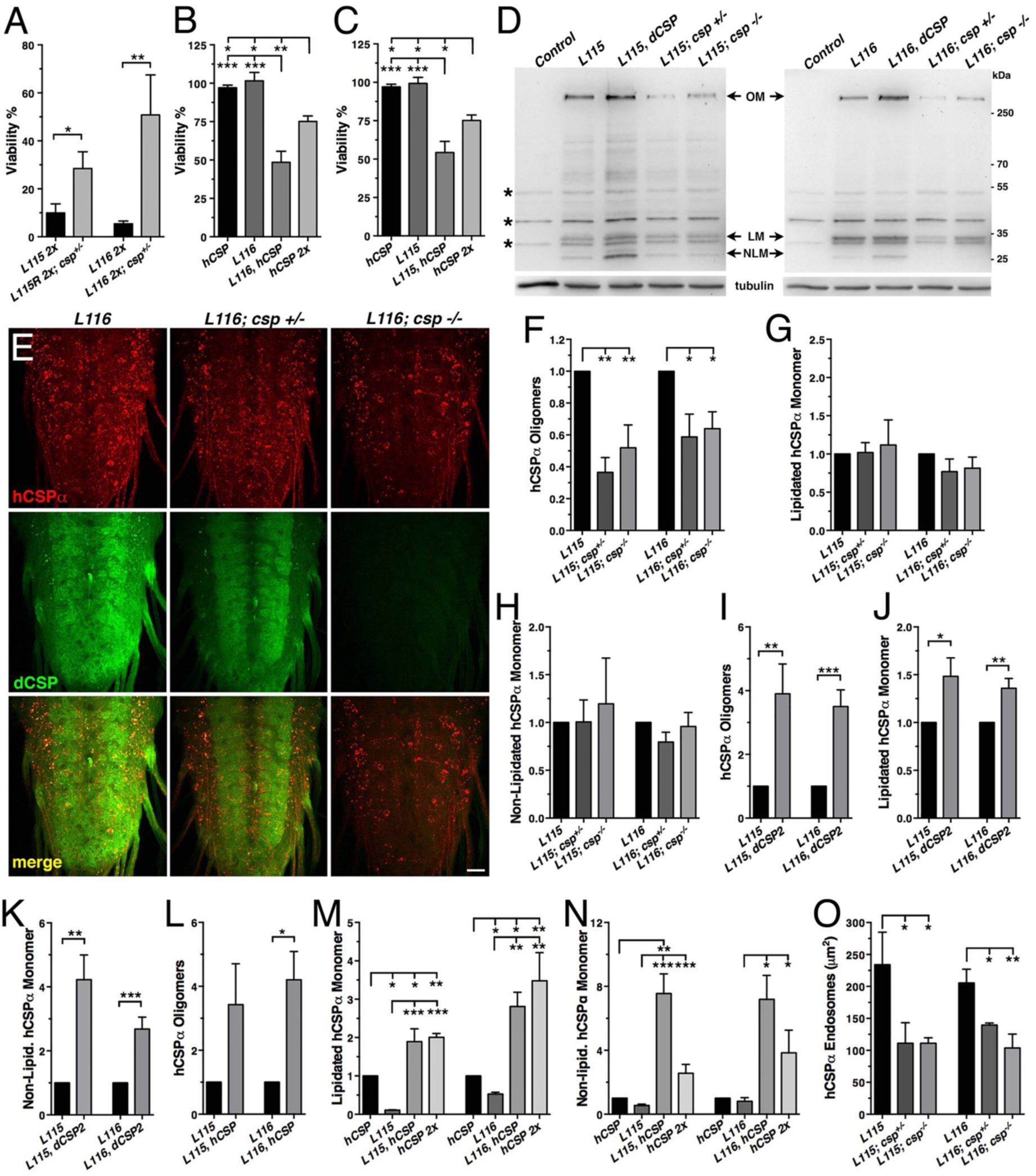
Altering wild type CSP levels modifies *CLN4* phenotypes. WT hCSPα, hCSP-L115 or - L116 were expressed in neurons from one or two transgenes (2x) with an elav driver in control (*w^1118^*), heterozygous *csp^X1/+^*, and homozygous *csp^X1/R1^* deletion mutants, or co-expressed with WT hCSPα or dCSP. Genotypes are indicated. **A-C.** Effects of reducing endogenous dCSP (**A**) or co-expressing WT hCSPα (**B-C**) on the viability of hCSP-L115 and -L116 mutant flies (N > 3; n > 144). **D.** Immunoblots of protein extracts from larval VNC probed for hCSPα and β-tubulin (loading control). hCSPα oligomers (OM), lipidated (LM), non-lipidated hCSPα monomers (NLM), and unspecific signals (*) are denoted. **E.** Larval VNC stained for hCSPα and dCSP. Scale bar, 20 μm. **F-N.** Effects of reduced (F-H) and increased endogenous dCSP (I-K) or WT hCSPα (L-N) levels on mutant hCSPα oligomers (F, I, L), lipidated monomers (G, J, M), and non-lipidated monomers (H, K, N). Signals were normalized to loading control and plotted as n-fold change of L115/L116 levels when expressed in a WT background (N = 5). **O.** Cumulative area of endosomal hCSPα accumulations in larval brains (N ≥ 3). Graphs display mean ± SEM. Statistical analysis used two-tailed unpaired *t* test (A, I-L), or one-way ANOVA (F-H, M-O); *, P < 0.05; **, P < 0.01; ***, P < 0.001.

Next, we determined whether altered WT CSP levels may also modulate the tendency of *CLN4* mutant hCSPα to form abnormal high-molecular weight oligomers. Reducing WT dCSP levels by expressing *CLN4* mutant hCSPα in hetero-or homozygous *dcsp* deletion mutants significantly attenuated the levels of SDS-resistant hCSP-L116 and -L115 oligomers (p<0.05; Fig. 7D, F). Levels of hCSP-L116 and -L115 oligomers were attenuated to a similar degree (p>0.5; Fig. 7F). Conversely, increasing WT dCSP levels by co-expressing a WT transgene together with a single *CLN4* mutant transgene increased the amount of hCSP-L115 and L116 oligomers (p<0.01, Fig. 7D, I). Coexpression of WT hCSPα also increased levels of hCSP-L116 and L115 oligomers (Fig. S5). However, only the increase in hCSP-L116 oligomers (p<0.5) was statistically significant (L115, p=0.09; Fig. 7L). In comparison, high-level overexpression of WT hCSPα from 2 transgenes did not induce significant protein aggregation (Fig. 2A, C, S5).

In contrast to the modulatory effects of WT CSP on mutant hCSPα oligomer levels, reducing or even abolishing WT dCSP levels had no effect on the levels of lipidated monomeric hCSP-L115 and - L116 (p>0.4; Fig. 7G). Coexpressing WT dCSP with *CLN4* mutant hCSPα slightly but significantly increased lipidated mutant hCSPα monomers (p<0.05; Fig. 7J) to levels that were similar to those of 2 copy WT hCSPα expression (Fig. 7M). This indicates that WT hCSPα may stabilize *CLN4* mutant hCSPα monomers.

Levels of non-lipidated monomeric *CLN4*-mutant hCSP were unaffected by reducing the gene dosage of endogenous WT dCSP (p>0.3; Fig. 7H). However, coexpression of WT dCSPα with *CLN4* mutant hCSPα increased levels of non-lipidated mutant hCSPα monomers (p<0.01; Fig. 7K). Similarly, coexpression of WT hCSPα also increased levels of non-lipidated hCSPα monomers (p<0.05; Fig.7N). This increase was non-additive since it was significantly larger than the increase induced by doubling WT hCSPα expression (p<0.05; Fig. 7N). Hence, increased levels of WT CSP may outcompete *CLN4* mutant hCSPα for palmitoylation or, alternatively, promote its depalmitoylation.

Finally, we tested whether altering levels of endogenous dCSP affects the endosomal accumulation of *CLN4* mutant hCSPα. Indeed, reducing dCSP levels by one gene copy significantly reduced the endosomal accumulation of both hCSP-L115 and -L116 in larval VNCs (p<0.05; Fig. 7E, O). Abolishing endogenous dCSP had a similar effect (Fig. 7O).

Taken together, the modulating effects of altering WT fly and human CSP levels on L115- and L116-induced phenotypes in viability, mutant hCSPα oligomerization and accumulation on endosomes suggest that the hCSP-L115 and -L116 alleles are hypermorphic gain of function mutations that increase an intrinsic activity. This conclusion is consistent with a recent study showing similar gene dosage effects of WT CSPα on hCSP-L115R phenotypes in mouse CSPα-deficient fibroblasts (Benitez and Sands, 2017). In addition, these findings uncover a strong correlation between the reduced viability of *CLN4* mutant flies and the degree of mutant hCSPα oligomerization and failure to be degraded.

### Partial loss of CSPα’s chaperone partner Hsc70-4 suppresses *CLN4*-induced phenotypes

The hypermorphic nature of *CLN4* mutations raised the possibility that at least some of the induced phenotypes are caused by an abnormally increased co-chaperone activity of CSP with the molecular chaperone Hsc70, which normally ensures efficient SV exo- and endocytosis by likely chaperoning SNARE proteins and Dynamin (Chandra et al., 2005; Nie et al., 1999; Sharma et al., 2012a; Sharma et al., 2011; Zhang et al., 2012). To address this possibility, we tested whether altered levels of SNARE proteins or Hsc70 may modify the eye phenotypes induced by GMR-driven expression of *CLN4* mutant hCSPα (Fig. 3).

Individual expression of Syntaxin 1A (Syx1A), neuronal Synaptobrevin (nSyb), or SNAP25 had no effect on the pigmentation and structure of the eye (not shown). However, co-expression of either Syx1A or nSyb with hCSP-L116 severely enhanced the pigmentation defect and significantly reduced the size of L116 mutant eyes (Fig. 8A). Overexpression or RNAi-mediated knockdown of SNAP25 had no effect (not shown), even though SNAP25 overexpression enhanced degenerative phenotypes of hCSPα knockout mice (Sharma et al., 2012a).

**Figure 8.**
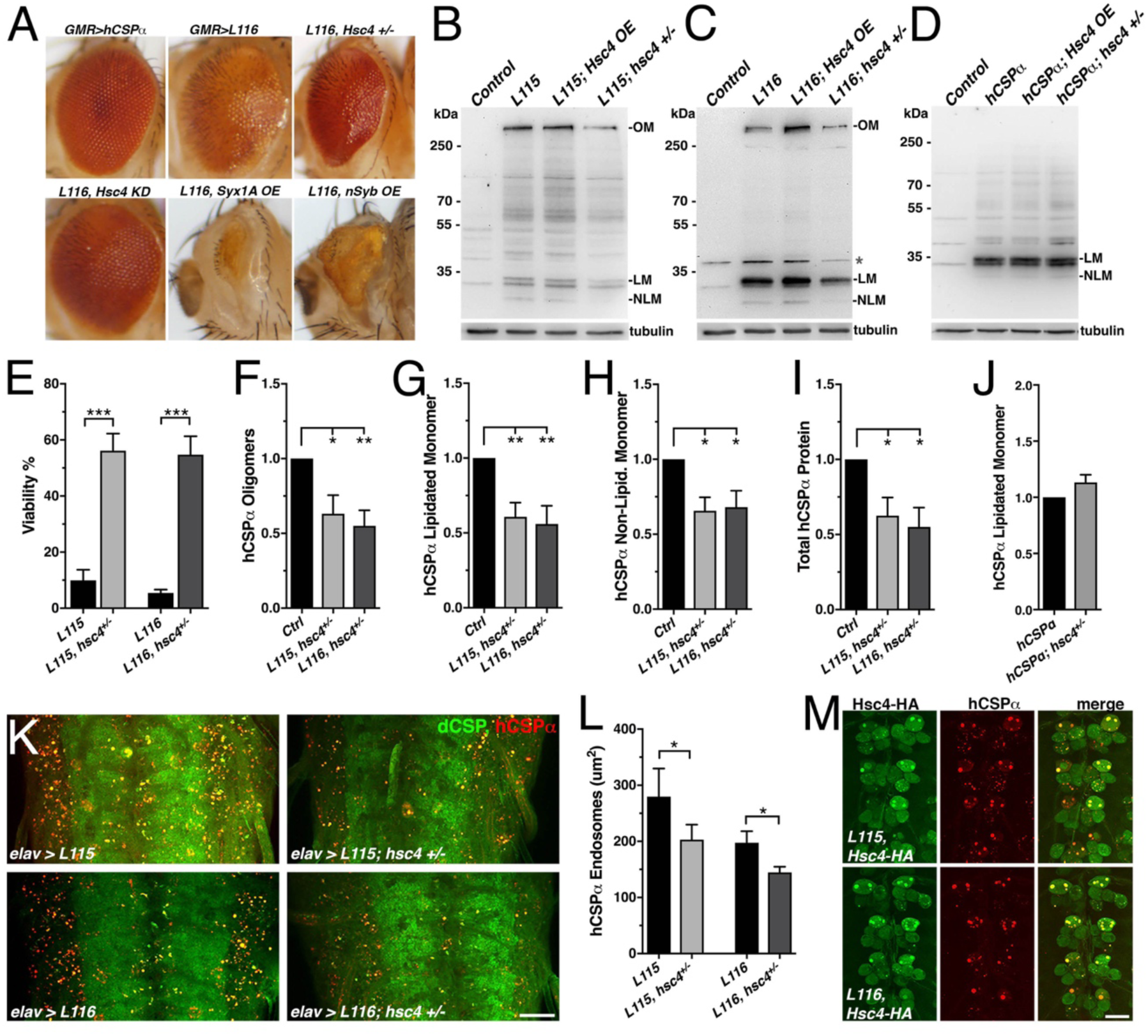
Reducing the gene dosage of Hsc4 attenuates *CLN4* phenotypes. **A.** Adult eyes of flies expressing WT hCSPα (control) or hCSP-L116 with a GMR-Gal4 driver in the absence or presence of a heterozygous Hsc4+/−(*hsc4^∆356^*) deletion, a Hsc4 KD, OE of Syx1A or Syb. **B-D.** Immunoblots of extracts form larval VNCs of indicated genotypes were probed for hCSPα. Lipidated monomeric hCSP (LM), non-lipidated hCSP (NLM) and hCSPα oligomers (OM) are indicated. Respective transgenes were expressed with elav-Gal; control was *w^1118^*. β-tubulin was used as loading control. **E.** Viability of animals expressing hCSPα-L115 or -L116 from two transgenes driven by elav-Gal4 in a control (*w^1118^*) or a heterozygous *hsc4^∆356^* deletion background (N ≥ 3; n > 74)**. F-I.** Effects of reduced *hsc4* gene dosage on elav-driven expression levels on mutant hCSPα oligomers (F), lipidated monomers (G), non-lipidated monomers (H) and total protein levels (I). Signals were normalized to loading control and plotted as n-fold change to levels of control (WT hCSPα; N > 6). **J.** Effect of reduced *hsc4* gene dosage on lipidated WT hCSPα monomer levels (N = 4). **K.** Segments of larval VNCs stained for dCSP and hCSPα. **L.** Cumulative area of mutant hCSPα accumulations in larval brains (N ≥ 4). **M.** Motor neuron somata of larval VNCs co-expressing HA-tagged Hsc4 with mutant hCSPα-L115 or -L116 stained for HA and hCSPα. Scale bars: 20 μm (K), 15 μm (M). Graphs display mean ± SEM. Statistical analysis used two-tailed unpaired *t* test (E, J), paired *t* test (L), or one-way ANOVA (F-I); *, P < 0.05; **, P < 0.01; ***, P < 0.001.

The genetic modifier effects of Syx1A and nSyb overexpression on *CLN4* eye phenotypes indicated that *CLN4* toxicity may be at least in part due to an altered chaperone activity of the CSP/Hsc70-complex. If so, one would expect that reducing levels of Hsc70 will suppress these phenotypes. This was indeed the case. Reducing endogenous Hsc70 levels by expressing hCSP-L116 in heterozygous Hsc70-4 (Hsc4) deletion mutants suppressed both the structural and depigmentation defects of L116 mutant eyes (Fig. 8A). Co-expression of three different UAS hairpin transgenes knocking down Hsc4 had a similar suppression effect (Fig. 8A and not shown). However, reducing the gene dosage of Hsc70-3 or Hsc70-5 had no effect (not shown), indicating that the suppression of L116-mutant eye phenotypes by reduced levels of Hsc4 is not a general effect of Hsc70 proteins.

To further test that Hsc4 may at least in part contribute to the toxicity of *CLN4* mutations, we examined effects of altered Hsc70 levels on the lethality induced by pan-neuronal expression of *CLN4* mutant hCSPα. Reducing the gene dosage of Hsc4 by expressing mutant hCSP-L115 or L116 in heterozygous *hsc4^∆356^* deletion mutants (Bronk et al., 2001) significantly suppressed their lethality (p<0.001; Fig. 8E), indicating that Hsc4 contributes to the toxicity of *CLN4* mutations.

Next, we examined to what degree a reduced gene dosage of Hsc4 may attenuate the oligomerization and endosomal accumulation of *CLN4* mutant hCSPα. Reducing Hsc4 protein levels significantly reduced the amount of hCSP-L115 and -L116 oligomers (p<0.05; Fig. 8B-C, F) but had no differential effects (p>0.8). Lowering Hsc4 levels also reduced the levels of lipidated and non-lipidated hCSP-L115 and -L116 monomers (p<0.05; Fig. 8G-H). Overall protein levels of mutant hCSP-L115 and - L116 were reduced (p<0.05; Fig. 8I). Reducing Hsc4 protein levels also suppressed the amount of *CLN4* mutant accumulations on endosomes in the larval brain (p<0.05; Fig. 8K-L, S6).

The overall reduction of *CLN4* mutant hCSPα could have been due to a general dependence of hCSPα expression on normal Hsc4 levels. However, this is unlikely the case since reducing Hsc4 levels had no effect on the levels of monomeric WT hCSPα (Fig. 8J), which was consistent with the normal dCSP levels observed in *hsc4* deletion mutants (Bronk et al., 2001). Hence, these findings suggest that Hsc4 significantly contributes to the pathology induced by *CLN4* mutations. Since Hsc4 colocalized with the abnormal accumulations of hCSP-L115 and -L116 (Fig. 8M), it may interfere with the prelysosomal trafficking and subsequent degradation of *CLN4* mutant hCSPα. This suggestion is consistent with the reduced overall mutant hCSPα levels that are induced by reduced Hsc4 levels (Fig. 8I).

## Discussion

To better understand pathological mechanisms underlying *CLN4*, we developed two *Drosophila* models of *CLN4* by expressing either *CLN4* mutant hCSPα or dCSP in neurons. Neuronal expression of either *CLN4* mutant hCSPα or dCSP mirrored key pathological features of post-mortem human *CLN4* brains, including reduced levels of palmitoylated hCSPα monomers and excessive formation of oligomers. Consistent with the idea that ubiquitination is likely a consequence of oligomerization, only ubiquitinated mutant hCSPα oligomers were detected but no ubiquitinated monomers. In general, the L115 mutation had stronger effects than L116, which has also been observed in post mortem brains and mammalian cell cultures (Diez-Ardanuy et al., 2017; Greaves et al., 2012; Henderson et al., 2016; Zhang and Chandra, 2014). Accordingly, expression of *CLN4* mutant hCSPα or dCSP in fly neurons provides a valid *CLN4* disease model.

### Dose-dependency of phenotypes in the *Drosophila CLN4* model

*CLN4* is a dominantly inherited disease with a variable post-juvenile onset between 25 to 46 years (Benitez et al., 2015; Bennett and Rakheja, 2013; Nijssen et al., 2003; Noskova et al., 2011). To model the dominant inheritance, we controlled the gene dosage of transgenically expressed *CLN4* mutant hCSPα by neuronally expressing either one or two mutant transgenes in the presence of two endogenous WT dCSP gene copies.

Single copy expression of *CLN4* mutant hCSPα did not cause detectable degeneration or lethality, even though significant levels of ubiquitinated hCSPα oligomers and abnormal prelysosomal accumulations of hCSPα were present. However, doubling the gene dosage of mutant hCSPα expression reduced the viability of *CLN4* mutant flies, which is likely due to neurodegeneration as indicated by fragmentated nuclear envelopes and “bloated” Golgi cisternae of larval neurons (Fig. 6B). The significant drop in viability indicated a sharp threshold of neurotoxicity, which was correlated with an increase in the levels of hCSPα oligomers and prelysosomal accumulations. Consistent with a neuronal failure, adult escapers showed reduced locomotor activity. A similar dose-dependent effect on toxicity was observed for the eye phenotype induced by GMR-driven expression of *CLN4* mutant hCSPα.

The dose-dependent build-up of pathological features towards a threshold of toxicity in the fly model is consistent with the late onset of *CLN4* in humans. It also conforms with a post-mortem report of a 37-year old *CNL4* patient in a clinically early pre-symptomatic stage who showed all of the pathological features of a terminal patient but had no significant signs of neurodegeneration or cognitive impairment (Benitez et al., 2015). Finally, the build-up of toxicity in the fly model agrees with the assumed general concept of late onset neurodegenerative diseases, which typically show a progressive accumulation of protein aggregates and defects in protein homeostasis over a long period that is met by an age-dependent declining capacity of protein homeostasis (Labbadia and Morimoto, 2015).

### *CLN4* mutations impair hCSPα’s synaptic localization

Since *CLN4* mutations reside in CSP’s palmitoylated CS domain, previous studies speculated that they may disrupt the trafficking and/or SV localization of hCSPα (Benitez et al., 2011; Greaves et al., 2012; Noskova et al., 2011). The *CLN4* fly model confirms this prediction in two ways: First, *CLN4* mutations severely reduced the synaptic levels of mutant hCSPα. Second, they caused abnormal accumulations of mutant hCSPα in axons and somata. Analogous *CLN4* mutations in fly dCSP had similar effects excluding the possibility that these defects are the consequence of expressing a human protein in fly neurons.

In principal, there are two possibilities how *CLN4* mutations may mislocalize CSP: 1) *CLN4* mutations may primarily impair palmitoylation of the CS domain, which is required for ER and Golgi exit, and for a stable association with SVPs and/or SVs (Greaves and Chamberlain, 2006; Greaves et al., 2008; Ohyama et al., 2007; Stowers and Isacoff, 2007). However, this possibility is unlikely the case since mutant hCSPα was not detectable in the ER or Golgi of fly neurons. In addition, levels of non-lipidated *CLN4* mutant hCSPα were similar to WT hCSPα when expressed from one transgenic copy. 2) *CLN4* mutations may primarily induce oligomerization by exaggerating the dimerization properties of the CS domain (Xu et al., 2010) such that self-association leads mostly to misfolded high-molecular weight oligomers that impair their association with SVPs or SVs. Notably, oligomerization of *CLN4* mutant hCSPα can be suppressed by alanine or leucine substitutions of a cluster of palmitoylated cysteines (C122-125) located next to the L115/116 mutations (Diez-Ardanuy et al., 2017). Since this suppression effect could not be explained by a loss of membrane association or reduced palmitoylation of the remaining cysteines in the CS domain (Diez-Ardanuy et al., 2017), it is consistent with the idea that *CLN4* mutations primarily induce oligomerization. Oligomerization is also the most plausible primary cause for the reduced synaptic levels of *CLN4* mutant hCSPα in fly neurons.

The ubiquitination of *CLN4* mutant hCSPα oligomers likely indicates that they are misfolded and thus targets for ubiquitination and degradation like other misfolded proteins (Wang et al., 2017). Consistently, only ubiquitinated mutant hCSPα oligomers but no ubiquitinated monomers were detected in fly neurons. Since WT dCSP (Fig. S4) and hCSPα are mostly degraded by lysosomes (Benitez and Sands, 2017; Chapel et al., 2013; Sambri et al., 2017; Schroder et al., 2007; Tharkeshwar et al., 2017), one expects that ubiquitinated mutant hCSPα oligomers are sorted into the endolysosomal pathway if they remain associated with membranes. This appears to be the case since the majority of abnormal mutant hCSPα accumulations in fly neurons represented prelysosomal endosomes, which were defined by the colocalization of ubiquitinated proteins and the prelysosomal markers LAMP1-GFP and HRS. Finally, the suppression of mutant hCSPα phenotypes by reduced dCSP or Hsc4 levels revealed a strong link between the oligomerization of mutant hCSPα, its re-routing onto prelysosomal membranes and its induced lethality (Figs. 7-8). Hence, these data suggest that *CLN4* mutations may primarily induce oligomerization of membrane-associated hCSPα, which causes ubiquitination and re-routing into lysosomal pathways.

The oligomerization and subsequent rerouting of mutant hCSPα is apparently triggered after exiting the Golgi, either during its association with SVPs, or after arriving at axon terminals and associating with SVs. The transition of SVP cargo onto SVs or endosomes is not well understood but in principle SVPs can fuse with SVs, the synaptic plasma membrane, and endosomes (Rizzoli, 2014). In contrast, SVs are linked to endosomes either by ultrafast endocytosis, or the ability of freshly endocytosed individual SVs to fuse with endosomes (Hoopmann et al., 2010; Soykan et al., 2017; Watanabe and Boucrot, 2017; Watanabe et al., 2013a; Watanabe et al., 2013b). Mutant hCSPα oligomers are potentially re-routed from both SVPs and SVs because some mutant hCSPα colocalizes with SVs at axon terminals. Nevertheless, re-routing of ubiquitinated oligomers is likely the main factor reducing synaptic levels of *CLN4* mutant hCSPα.

### *CLN4* mutations impair prelysosomal trafficking of hCSPα

Axonal prelysosomal endosomes containing mutant hCSPα are likely retrogradely trafficked for lysosomal processing since degradative lysosomes containing cathepsin B/D are essentially absent from distal axons (Cai et al., 2010; Cheng et al., 2018; Gowrishankar et al., 2015). Both the axonal trafficking as well as the maturation of prelysosomal endosomes containing mutant hCSPα is potentially impaired. A physical defect in axonal trafficking is indicated by the large size of the hCSPα accumulations and by the frequent occurrence of large and abnormal EM-dense membrane structures in axons of *CLN4* mutants, which have the potential to “clog” axons.

A reduced efficiency in the maturation of mutant hCSPα-positive prelysosomal endosomes is indicated by their abnormally large size and by their failure to co-label with the PI_3_P lipid reporter FYVE-GFP. The latter may be due to two alternatives: First, hCSPα-positive endosomes may have normal PI_3_P levels but most of it is bound by PI_3_P-binding proteins. At least in part, this could be facilitated by HRS (Raiborg et al., 2001b), which strongly co-labels with hCSPα accumulations. Alternatively, PI_3_P could be depleted from these endosomal membranes, which then would imply that HRS is abnormally retained. Since only a small number of mutant hCSPα accumulations co-localized with the early endosome marker Rab5 but essentially no accumulations colocalized with the late endosomal or lysosomal markers Rab7 and spinster, it is possible that mutant hCSPα may impair the transition from early to late endosomes.

Finally, the frequent occurrence of abnormal, multi-laminar structures containing EM-dense membranes and abnormal autophagosome-like structures indicates that *CLN4* mutations impair steps of membrane trafficking in axons and somata. Since similar ultrastructural defects were seen after disruption of ESCRT function (Doyotte et al., 2005; Lee et al., 2007), the observed abnormal structures of *CLN4* mutants may indicate a defect in prelysosomal trafficking, which is consistent with the accumulation of mutant hCSPα on prelysosomal membranes including ATG8/LC3-GFP positive autophagosomes. Alternatively, at least some of the abnormal membrane structures of CLN4 mutants could be the consequence of defects in the unconventional secretion of misfolded cytosolic proteins, which is facilitated by CSPα at least in non-neuronal cells (Xu et al., 2018).

### *CLN4* mutations increase overall levels of ubiquitinated proteins

Next to an impaired localization and prelysosomal processing of mutant hCSPα, the *CLN4* fly model also provides evidence for a protein homeostasis defect. Both of the *CLN4* mutations caused an increase in protein ubiquitination in axons and neuronal somata that was not associated with accumulations of *CLN4* mutant hCSPα. This excessive ubiquitination may directly arise from a reduced CSP/Hsc70 chaperone function on SVs since *CLN4* mutant CSPα’s ability to stimulate the ATPase activity of Hsc70 progressively deteriorates *in vitro* (Zhang and Chandra, 2014). Alternatively, the excessive ubiquitination may be a secondary consequence of defects induced by *CLN4* mutations. Both cases are consistent with the increased expression of proteins associated with proteasomal degradation and increased proteasomal activity in response to a loss of CSP’s chaperone function in CSPα KO mice (McCue et al., 2015; Sharma et al., 2012b). In addition, significant proteomic and lipidomic changes were found in *CLN4* mutant post-mortem brains (Henderson et al., 2016).

### *CLN4* alleles genetically resemble hypermorphic gain of function mutations

Several hypotheses regarding the genetic nature of the dominant *CLN4* alleles L115 and L116 have been suggested. Based on CSPα’s neuroprotective role (Fernandez-Chacon et al., 2004; Zinsmaier et al., 1994), it has been suggested that the progressive oligomerization of mutant hCSPα may cause a dominant-negative effect by sequestering WT CSPα (Greaves et al., 2012; Henderson et al., 2016; Noskova et al., 2011; Zhang and Chandra, 2014). However, *CLN4* mutant hCSPα had no effect on the synaptic and overall levels of endogenous WT dCSP, which is inconsistent with the idea of sequestering WT CSP. In addition, genetic characterization of the *CLN4* alleles revealed a complex dependence of *CLN4* phenotypes on WT CSP levels, which had either a positive or no effect. Both exclude the possibility that the alleles L115 and L116 exert dominant-negative effects.

Reducing the gene dosage of WT dCSP had no effect on the levels of monomeric palmitoylated mutant hCSPα, but suppressed the lethality, oligomerization and endosomal accumulation of mutant hCSPα. *Vice versa*, increasing WT dCSP or hCSPα levels enhanced the latter phenotypes. Hence, these modulatory effects of WT CSP on *CLN4* phenotypes suggest that both *CLN4* alleles act genetically as hypermorphic gain of function mutations increasing an intrinsic function of the protein. Notably, a similar dependence of *CLN4* phenotypes on WT CSPα was observed in primary fibroblasts of CSPα-deficient mice after co-expressing WT CSPα with CSPα-L115R, which increased levels of CSPα oligomers and lysotracker signals (Benitez and Sands, 2017). Hence, *CLN4* mutations cause complex genetic effects that vary among the induced phenotypes, as speculated earlier (Greaves et al., 2012; Henderson et al., 2016; Zhang and Chandra, 2014; Zhang et al., 2012).

The dependence of *CLN4* induced pathology/toxicity on WT CSP function paralleled the modulatory effects of hCSPα’s main interaction partner, Hsc70. Similar to reducing WT dCSP levels, reducing levels of Hsc70-4 by one gene copy suppressed the lethality and eye phenotypes induced by *CLN4* mutant hCSPα, its oligomerization and endosomal accumulation. These modulatory effects are driven either by an individual activity of Hsc70 itself, or by a synergistic activity of the CSP/Hsc70 complex. Neither of these activities are necessarily localized to SVs. In support of the latter, co-expressed HA-tagged Hsc70-4 co-localized with the accumulations of *CLN4* mutant hCSPα on endosomes, which may interfere with the dissociation of clathrin coats that are anchored by the ESCRT component HRS (Raiborg et al., 2001a). Alternatively, the respective Hsc70 activity may be associated with endosomal microautophagy (Sahu et al., 2011; Uytterhoeven et al., 2015) or the unconventional secretion of misfolded cytosolic proteins (Xu et al., 2018). At least for this fly model, an activity of Hsc70 associated with chaperone-mediated autophagy (Kaushik and Cuervo, 2012) can be excluded since the critical lysosomal translocator LAMP-2 is absent in flies (Uytterhoeven et al., 2015).

### Concluding Remarks

This initial study of the fly model revealed a number of novel and important insights into the pathology of *CLN4*, most notably, the unexpected genetic nature of *CLN4* mutations, the mislocalization of mutant hCSPα, a general protein homeostasis defect, and a link between *CLN4* mutations and prelysosomal processing defects. The modulating effects of altered WT CSP or Hsc70 levels on key *CLN4* phenotypes also provided a strong correlation between the premature lethality of *CLN4* mutant flies, oligomerization and accumulation of mutant hCSPα on prelysosomal endosomes. Accordingly, the neurotoxicity of *CLN4* mutant hCSPα may be at least in part due to the formation of ubiquitinated hCSPα oligomers that then progressively accumulate on prelysosomal endosomes and interfere with their processing and potentially their retrograde axonal trafficking.

The pathological mechanisms underlying *CLN4* appear to be different from those of most other NCLs, which are typically due to loss of function mutations in genes that mediate lysosomal function, ER-lysosomal trafficking, or protein lipidation (Carcel-Trullols et al., 2015; Cooper et al., 2015; Cotman et al., 2013; Mole and Cotman, 2015; Warrier et al., 2013). As such, the pathological mechanisms underlying *CLN4* appear similar to those of “classical” neurodegenerative diseases that are induced by a progressive build-up of protein oligomers/aggregates that then leads to a failure of protein homeostasis and/or lysosomal pathways (Abramov et al., 2009; Labbadia and Morimoto, 2015; Lansbury and Lashuel, 2006; Muchowski and Wacker, 2005; Neefjes and van der Kant, 2014). While we are just beginning to understand the complicated nature of *CLN4*, the fly model provides a valuable tool for future work to dissect pathological mechanisms underlying *CLN4*.

## Material and Methods

### *Drosophila* Strains and Husbandry

All flies were raised at 23°C on standard cornmeal culture media with a 12/12 light-dark cycle unless otherwise specified. Gal4 driver strains (elav-Gal4 (C155), nSyb-Gal4, GMR-Gal4) and UAS strains expressing EGFP-nSyb, GFP-myc-2xFYVE, TSG101 RNAi (*P[y^+t7.7^, v^+t1.8^=TRiP.GLV21075]attP2*), hLAMP1-GFP, nSyb-EGFP, Rab5-YFP, Rab7-YFP were obtained from the Bloomington *Drosophila* Stock Center (BDSC, Bloomington, Indiana). A UAS strain expressing Venus-Rab7 was obtained from R. Hiesinger (Freie Universität, Berlin, Germany). Hsc70-4 deletion (Δ356) and UAS Hsc70-4 lines were generated previously by us (Bronk et al., 2001). Fly strains containing UAS-transgenes expressing HA-Hsc70-4 (Uytterhoeven et al., 2015) were obtained from P. Verstreken (VIB Center for the Biology of Disease, Leuven, Belgium).

### Generation of UAS Transgenes

To generate transgenes that individually express human WT, L115R- and L116Δ-mutant CSPα under the transcriptional control of the Gal4/UAS system (Brand and Perrimon, 1993), the open reading frames of modified rat cDNAs encoding human WT and mutant CSPα (Zhang and Chandra, 2014) were PCR amplified using a forward primer containing a NotI restriction site and a *Drosophila* consensus Kozak sequence (5’ GAGCGGCCGCCAAAATGGCTGACCAGAGGCAGCGCTC 3’) and a reverse primer containing a KpnI site just after the stop codon (5’ CATGGTACCTTAGTTGAACCCGTCGGTGT GATAGCTGG 3’). The obtained PCR products were cleaved with NotI and KpnI and directionally cloned into a NotI/KpnI cleaved pBID-UASC vector (Wang et al., 2012).

cDNAs encoding WT and *CLN4* mutant dCSP were synthesized *de novo* (GenScript, Piscataway, NJ) using cDNA sequences encoding the open reading frame of CSP2 (CSP-PC, accessionID:NP_730714) as template (Nie et al., 1999). The mutations V117R and I118Δ were introduced into dCSP2 to generate *CLN4* mutant dCSP. Both V117R and I118Δ are analogous to the human *CLN4* mutations L115R and L116Δ, respectively. The synthesized cDNAs contained a NotI site followed by a Kozak sequence at the 5’ end and a Kpn1 site after the stop codon at the 3’ end and were inserted into pUC57 plasmids. For transgene expression, the cDNAs were then directionally subcloned into a pBID-UASC vector using the NotI and KpnI cleavage sites.

Transgenic animals were generated by ϕC31-based integration (Bischof et al., 2007) of the UAS transgenes into the attP site at 22A2 on chromosome 2 (2L:1476459..1476459). pBID-UASC-CSP-x plasmids were injected into *y^1^ M[vas-int.Dm]ZH-22A w*; M[3xP3-RFP.attP’]ZH-22A* (BDSC #24481) embryos (Rainbow Transgenic Flies, Camarillo, CA). At least two independent recombinant strains were obtained for each transgene and out-crossed to exchange non-recombinant chromosomes. The *3xP3-RFP* cassette was removed by loxP-mediated recombination as described (Bischof et al., 2007). Homozygous strains containing UAS transgenes were established in a genetic background representing WT control (*w^1118^*) and *dcsp* deletion mutants (*w^1118^; dcsp^X1^*).

Crosses for single copy transgene expression were made by crossing homozygous male *w^1118^, elav-Gal4[C155]* flies to homozygous *w^1118^; M[UAS::CSP-x.]ZH-22A* female flies yielding female F1 progeny expressing CSP-x (*w^1118^, elav-Gal4[C155]/w^1118^; M[UAS::CSP-x]ZH-22A /+*) and male progeny containing a silent (non-expressed) transgene (*w^1118^; M[UAS::CSP-x]ZH-22A/+*). Crosses for 2 copy expression were made by crossing females *w^1118^, elav-Gal4[C155]; M[UAS::CSP-x]ZH-22A/CyO, Actin-GFP* to male *w^1118^; M[UAS::CSP-x]ZH-22A*. Female F1 progeny heterozygous for the *elav* driver and homozygous for the UAS transgene (*w^1118^*, *elav-Gal4[C155]/w^1118^; M[UAS::CSP-x}ZH-22A*) were selected for analysis.

### dCSP null rescue

All genetic rescue experiments employing a *csp^−/−^* deletion null genetic background expressed WT or *CLN4* mutant hCSPα from a single transgene. Female *w^1118^, elav-Gal4[C155]; csp^R1^/TM6 Tb Sb* flies were crossed to *w^1118^; M[UAS::hCSP-x]ZH-22A; csp^X1^/TM6 Tb Sb* males or *w^1118^; csp^X1^/TM6 Tb Sb* males for control. Male progeny (*w^1118^, elav-Gal4[C155]; M[UAS::hCSP]ZH-22A/+; csp^X1/R1^*) were used because of higher expression and better rescue. 10-15 freshly enclosed flies of the respective genotypes were cultured in separate vials, counted and transferred to fresh food every 2 days.

### Viability

Flies were raised at 27-28°C to enhance Gal4-dependent transgene expression. The following crossing scheme was used: For single copy expression, hemizygous *w^1118^, elav-Gal4[C155]* males were crossed to homozygous *w^1118^; UAS::CSP-x* females yielding non-expressing F1 control males (*w^1118^; UAS::CSP-x/+*) and CSP-expressing F1 females (*w^1118^, elav-Gal4[C155]/w^1118^; UAS::CSP-x/+*). For 2 copy expression, hemizygous *w^1118^, elav-Gal4[C155]; UAS::CSP-x/CyO, Actin-GFP* males were crossed to homozygous *w^1118^; UAS::CSP-x* females yielding non-expressing F1 males (*w^1118^; UAS::CSP-x*) and CSP-expressing F1 females (*w^1118^, elav-Gal4[C155]/w^1118^; UAS::CSP*). Freshly enclosed flies of the respective genotypes were cultured in separate vials, counted and transferred to fresh food every 2 days. Viability was determined by taking the ratio of females and male flies.

### Immunostainings

Wandering 3^rd^ instar larvae were dissected in Sylgard-coated dishes containing cold Ca^2+^ free HL3 solution (in mM: 70 NaCl, 5 KCl, 20 MgCl_2_, 10 NaHCO_3_, 5 Trehalose, 115 sucrose, 5 HEPES, pH 7.2). Dissected larvae were fixed for 20 (VNC) or 45 minutes (NMJs) in 4% formaldehyde solution (Electron Microscopy Sciences, Hatfield, PA) in phosphate-buffered saline (PBS), pH 7.3 at room temperature (RT). PBS (pH 7.3) supplemented with 0.2% Triton X-100 (PBST) was used for immunostainings of larval NMJs while PBS (pH 7.3) supplemented with 0.4% Triton X-100 was used for immunostainings of larval VNCs. After washing 3 times for 10 minutes in PBST at RT, larvae were incubated with primary antibodies diluted in PBST overnight at 4°C, washed 3x for 15 minutes at RT, incubated with secondary antibodies diluted in PBST for 2 hours at RT or overnight at 4°C and finally washed 3x with PBST for 15 minutes at RT. Confocal images were acquired the same day, otherwise preparations were post-fixed. The following antibodies and dilutions were used: rabbit anti-CSPα, 1:2,000 (Enzo Life Sciences Cat# VAP-SV003E, RRID:AB_2095057); mouse anti-dCSP, 1:250 (DSHB Cat# DCSP-1 (ab49), RRID:AB_2307345, (Zinsmaier et al., 1990)); mouse anti-GFP, 1:1,000 (DSHB Cat# DSHB-GFP-12A6, RRID:AB_2617417); guinea pig anti-HRS, 1:2,000 ((Lloyd et al., 2002), H. Bellen, Baylor College of Medicine, Houston, TX); rat anti-HA, 1:200 (Roche Cat# 3F10, RRID:AB_2314622); rabbit anti-GM130, 1:200 (Abcam Cat# ab31561, RRID:AB_2115328); goat anti-Golgin245, 1:2000 (DSHB Cat# Golgin245, RRID:AB_2618260), goat anti-GMAP, 1:2000 (DSHB Cat# GMAP, RRID:AB_2618259); mouse anti Rab7, 1:100 (DSHB Cat# Rab7, RRID:AB_2722471, (Riedel et al., 2016)); mouse anti-Syx1A, 1:200 (DSHB Cat# 8c3, RRID:AB_528484); mouse anti-ubiquitin-conjugated protein, 1:2000 (Enzo Life Sciences Cat# BML-PW8810, RRID:AB_10541840); goat anti-HRP Alexa Fluor 647-conjugated, 1:500 (Jackson ImmunoResearch Labs Cat# 123-605-021, RRID:AB_2338967); goat anti-mouse IgG1 Alexa Fluor 488-conjugated, 1:500 (Thermo Fisher Scientific Cat# A-21121, RRID:AB_2535764); donkey anti-rabbit IgG (H+L) Cy3-conjugated, 1:500 (Jackson ImmunoResearch Labs Cat# 711-165-152, RRID:AB_2307443); goat anti-guinea pig IgG (H+L) Alexa Fluor 488-conjugated, 1:1000 (Thermo Fisher Scientific Cat# A-11073, RRID:AB_2534117); goat anti-rat IgG (H+L) Alexa Fluor 488-conjugated, 1:500 (Jackson ImmunoResearch Labs Cat# 112-545-167, RRID:AB_2338362).

### Confocal Imaging

Stained preparations were imaged with an Olympus microscope BX50WI equipped with a confocal laser scanner (FluoView 300), a 60X water-immersion objective (LUMPLFL; N.A., 0.9), a multi-argon (630), a green HeNe (430), and a red HeNe (630) laser. Optical sections in the vertical axis were acquired using 0.8 μm (NMJs) or 1.5 μm (VNC) intervals for optical sectioning using Fluoview software. Images were analyzed offline using ImageJ software (FIJI v1.50, NIH).

For quantification of fluorescence signals, control and mutant larvae were dissected in the same dish such that fixation and antibody incubation were performed identically. All samples were imaged with the same laser settings. Fluorescence intensity per area was determined from a region of interest (ROI) encompassing single synaptic boutons by using Image J Software.

For quantification of CSP accumulations in VNC, z-projections with maximal intensity of optical sections were generated were cropped to a defined volume of 60 × 80 × 45 μm (216,000 μm^3^). These image stacks represented the dorsal-most part of hemi-segments A4 and A5. After thresholding for background subtraction, ROIs were drawn around all visible CSP-positive punctae; for those appearing in multiple optical sections only the section with the brightest signal was used. ROI areas were compiled to compare the cumulative area among genotypes.

### Western Blot Analysis

Larval brains were dissected from wandering 3^rd^ instar larvae in in phosphate-buffered saline (PBS, pH 7.3) and 5 brains transferred to 60 µL buffer (2% SDS, 10% Glycerol, 60 mM Tris pH 6.8, 0.005% Bromophenol Blue, and 100 mM DTT). Brains were homogenized with a p200 pipette tip, boiled for 5 minutes and centrifuged for 3 minutes at 2000 g. The soluble fraction was transferred to a fresh tube, boiled for 1 min and an equivalent of ∼1.5 brains was immediately loaded onto a 10% acrylamide gel for SDS-PAGE at 80V (Mini-Protean Cell, BioRad, Hercules, CA). Separated proteins were blotted onto nitrocellulose membranes at 20 V for 6 minutes using an iBlot system (Invitrogen, Carlsbad, CA). After transfer, the blot was blocked for 30 minutes using 1% bovine serum albumin fraction V (#BP1600-100, Fisher Scientific) in 0.2% PBS supplemented with Tween-20, pH 7.3 (PBST). Blots were incubated with primary antibodies overnight at 4°C, washed, and incubated with HRP-conjugated secondary antibodies for 2 hours at 4°C. To normalize for protein loading, blots were stripped (#21059; ThermoFisher Scientific) for 15 minutes at RT and immunostained for β-tubulin. Blots were imaged using a BioRad Western Clarity ECL kit and ChemiDoc XRS imaging system. Protein band intensities were quantified via densitometry analysis with Quantity One software (Bio-Rad). Raw intensity values were normalized to the measurement of a β-tubulin loading control and expressed as the n-fold change to an appropriate control genotype. Antibodies were diluted in PBST and used at the following concentrations: rabbit anti-CSPα 1:20,000 (Enzo Life Sciences Cat# VAP-SV003E, RRID:AB_2095057); mouse anti-dCSP, 1:20 (DSHB Cat# DCSP-1 (ab49), RRID:AB_2307345, (Zinsmaier et al., 1990)); mouse anti-ubiquitin-conjugated protein, 1:2000 (Enzo Life Sciences Cat# BML-PW8810, RRID:AB_10541840); mouse anti-β-tubulin, 1:1000 (DSHB Cat# E7, RRID:AB_528499); goat anti-mouse IgG HRP-conjugated, 1:5000 (Thermo Fisher Scientific Cat# 32430, RRID:AB_1185566; goat anti-rabbit IgG (H+L) HRP-conjugated 1:10,000 (Thermo Fisher Scientific Cat# A16096, RRID:AB_2534770).

### Hydroxylamine Treatment

Adult heads were homogenized in 2% SDS, 100 mM DTT, 60 mM Tris pH 6.8, and then treated with 0.5M hydroxylamine pH 7.0 or 0.5 M Tris pH 7.0 (final concentration) for 24 hours at RT before being diluted 3-fold using 1x Laemmli buffer. Samples were boiled for 5 minutes before use for SDS-PAGE.

### Immunoprecipitation

Young adult flies (1-3 day post-eclosion) were flash frozen in liquid N_2_ and stored at −80°C. Heads were enriched by brushing frozen heads on double metal sieves under liquid N_2_. Approximately 100 μL of heads were homogenized in 500 μL IP buffer containing: 1% Triton X-100, 30 mM Tris-HCl (pH 7.2), 150 mM KCl, 0.5 mM MgCl_2_. Homogenates were repeatedly centrifuged at 22,000 rpm to remove insoluble debris. The cleared supernatant was incubated with rabbit anti-CSPα at 1:200 (f.c.) for 1 hour on a rotator at 4°C and then incubated with 50 μL protein A/G PLUS-Agarose (#sc-2003, Santa Cruz Biotechnology, Dallas, Texas) for 2 hours at 4°C. Agarose beads were spun down with a tabletop centrifuge at 2000g and the supernatant was removed and stored. The beads were washed 4x with IP buffer by repeated resuspension and centrifugation. Beads were finally resuspended in Laemmli buffer, boiled for 5 minutes, centrifuged for 2 minutes and transferred to a fresh tube for SDS-PAGE.

### Electron Microscopy

Larvae were rapidly dissected in 4% PFA and then fixed overnight in fresh fixative containing: 3% glutaraldehyde, 1.5% formaldehyde, 2 mM CaCl_2_, 0.1M sodium cacodylate, pH 7.3. Samples were rinsed in 0.1M cacodylate before being fixed with 2% OsO4 for 2 hours. Next, samples were washed with HPLC grade H_2_O before being dehydrated with ethanol in a stepped series (30%, 50%, 70%, 90%, 95%,100%) followed by 100% acetone. Tissue was infiltrated with Durcupan ACM plastic embedding media (#14040, Electron Microscopy Sciences, Hatfield, PA) through progressive mixtures with acetone (25%, 50%, 75%, 100%), hardened at 60°C, trimmed and sectioned with a diamond knife (70 nm). Sections were poststained with 4% uranyl acetate and lead citrate. Images were obtained on a JEOL 1200EX with an AMT XR80M-B camera running AMT software. For publication, figures were compiled and prepared with Photoshop CC (Adobe). Contrast and intensity of images was minimally adjusted. Images were cropped as needed.

### Data and Statistical Analysis

Data from at least 3 independent animals or experimental trials were used for statistical analysis. Data are represented as mean, and error bars represent SEM. Gaussian distribution of data was assessed using a D’Agostino & Pearson omnibus or Shapiro-Wilk normality test using Prism software (GraphPad Software). Statistical significance was assessed by either a two-tailed *t* test, Mann-Whitney, one-way ANOVA (Kruskal-Wallis for non-parametric data) test with appropriate post-hoc tests using Prism software). P values <0.05, <0.01, and <0.001 are indicated in graphs with one, two, and three asterisks, respectively.

## Acknowledgements

We thank Drs. Hugo J. Bellen (Baylor College of Medicine, Houston TX, USA), P. Robin Heisinger (Freie Universität, Berlin, Germany), Patrik Verstreken (VIB-KU Leuven Center for Brain & Disease Research, Leuven, Belgium), and the Developmental Studies Hybridoma Bank at the University of Iowa for antibodies and/or fly strains. We thank Patty Jansma, Andrew Wellington, Stephan Dong, Marija Zaruba, Mays Imad, and Milos Babic for their technical help and critical feedback. This work was supported by grants from NINDS (R01 NS083849 to SSC (PI) and KEZ (subaward); R21 NS094809 to KEZ).

The authors declare no competing financial interests.

## Author Contributions

E. Imler, S.S. Chandra and K.E. Zinsmaier conceived the study. E. Imler generated the transgenic strains and performed the majority of the experiments. J.S. Pyon and S. Kindelay helped with data collection. Y. Zhang and S.S. Chandra provided reagents. E. Imler and K.E. Zinsmaier analyzed and interpreted data. E. Imler and K.E. Zinsmaier designed experiments and wrote the manuscript.

