## Supplemental Material for "A *Drosophila* model of neuronal ceroid lipofuscinosis *CLN4* reveals a hypermorphic gain of function mechanism"

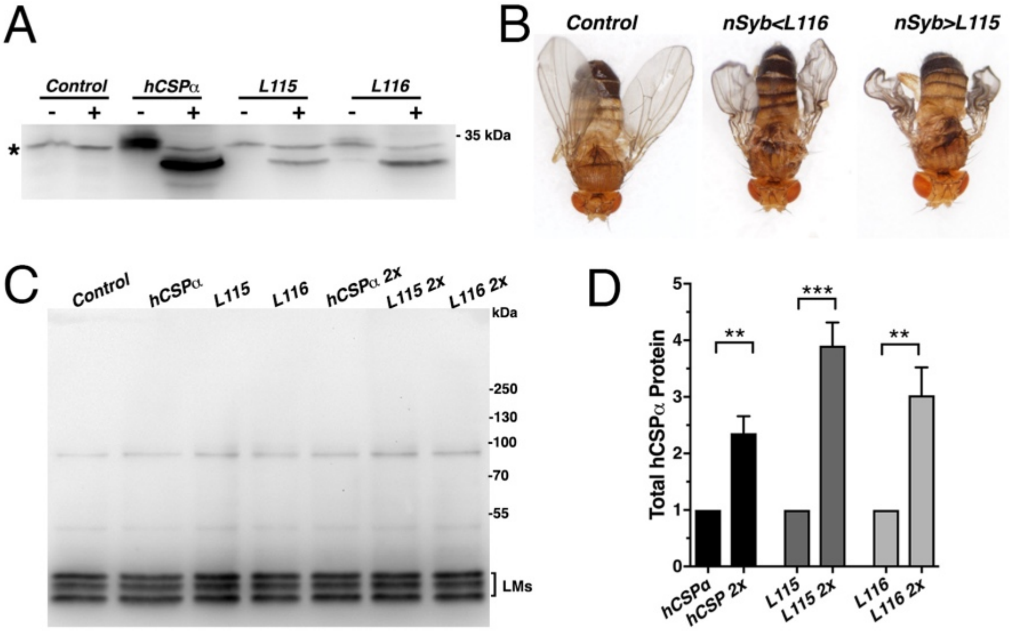


**Figure S1. Phenotypic effects of *CLN4* mutations. A.** Lipidation of WT hCSPα, hCSP-L115 and -L116. Immunoblot of larval VNC extracts from control ($w^{1118}$) and animals expressing elav-driven hCSPα, hCSP-L115 or -L116 were probed for hCSPα. Samples were treated overnight with either 0.5 M hydroxylamine (+) or equimolar Tris (-). Asterisk denotes unspecific signal. **B.** Adult control ($w^{1118}$) and animals expressing nSyb-driven hCSP-L115 or -L116 in neurons. **C.** Immunoblot probed for dCSP of larval brain extracts from control ($w^{1118}$) and animals expressing WT hCSPα, hCSPα-L115 or -L116 from either one or two (2x) transgenes with an elav driver. The lipidated monomeric dCSP isoforms (LMs) are indicated. **D.** Increase in total protein levels of WT hCSPα, hCSP-L115 and -L116 induced by expressing either one or two transgenes (2x). Signals were normalized to loading control and plotted as n-fold change to respective expression levels from one transgene (mean ± SEM; N = 6, two-tailed unpaired *t* test; **, $P < 0.01$; ***, $P < 0.001$.

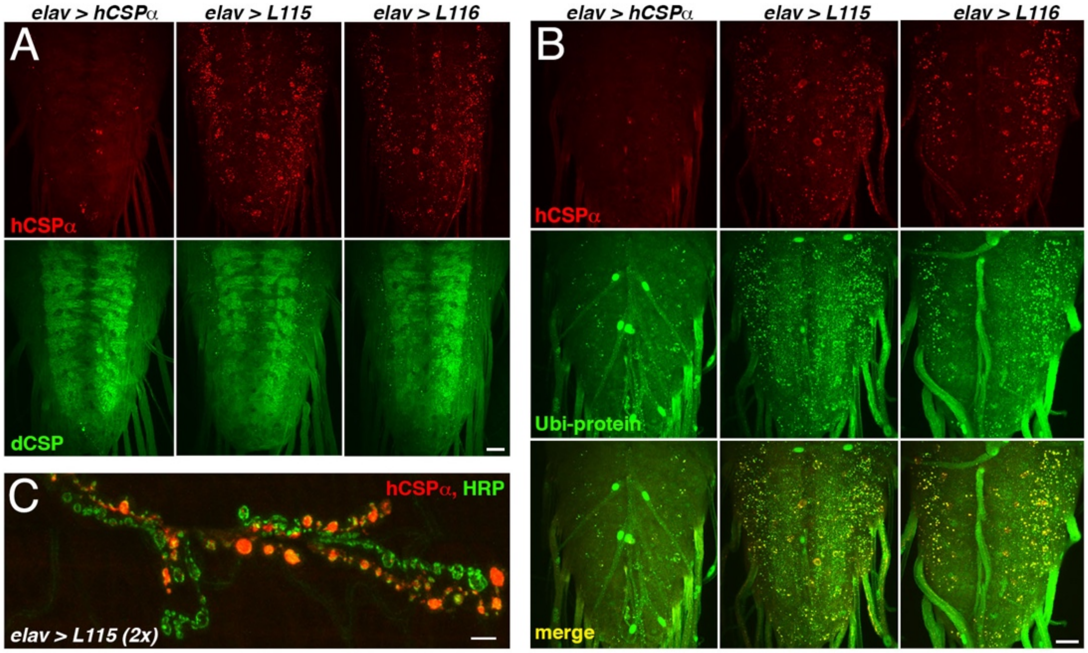


**Figure 2. hCSP-L115 and -L116 abnormally accumulate with endogenous dCSP and ubiquitinated proteins**. WT and mutant hCSPα (L115 and L116) were expressed in larval neurons with an elav-Gal4 driver from 1 (A-B) or two (C) transgenes (2x). **A-B.** Larval VNCs of indicated genotypes immunostained for hCSPα and dCSP (A) or lysine-linked-ubiquitin (B). **C.** Larval NMJ stained for hCSPα and HRP showed occasionally extreme accumulations of hCSP-L115 in synaptic boutons. Scale bar, 20 μm (A-B), 5 μm (C).

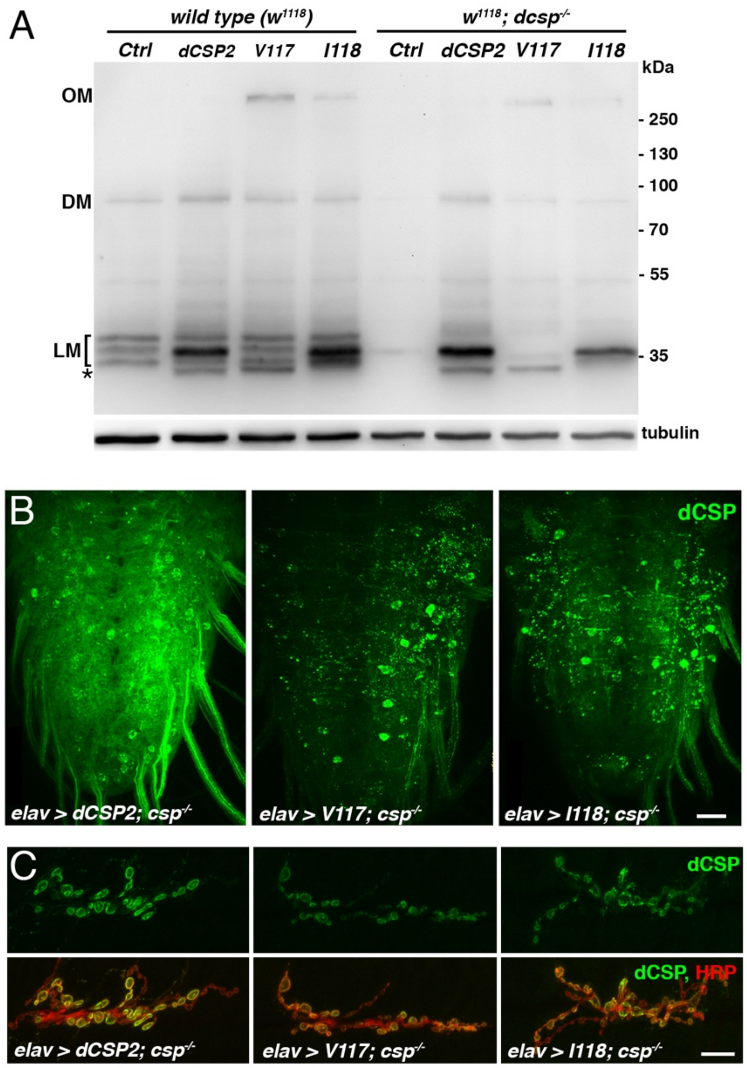


**Figure S3. Effects of *CLN4* analogous mutations in dCSP.** WT dCSP2, dCSP-V117R (analogous to human L115R) and dCSP-I118Δ (analogous to human L116Δ) were expressed in larval neurons with an elav-Gal4 driver from one transgene in otherwise wild type control ($w^{1118}$) or homozygous *dcsp* null mutants ($dcsp^{X1/R}$). **A.** Immunoblot of larval VNC extracts of indicated genotypes probed for dCSP. Signals corresponding to lipidated dCSP monomers (LM), dimers (DM), and high-molecular weight oligomers (OM) are indicated. Asterisk denotes partially lipidated dCSP. β-tubulin was used as loading control. **B.** Larval VNCs of indicated genotypes immunostained for dCSP. **C.** Larval NMJs of indicated genotypes stained for dCSP and HRP. Scale bar, 20 µm (A), 10 µm (C).

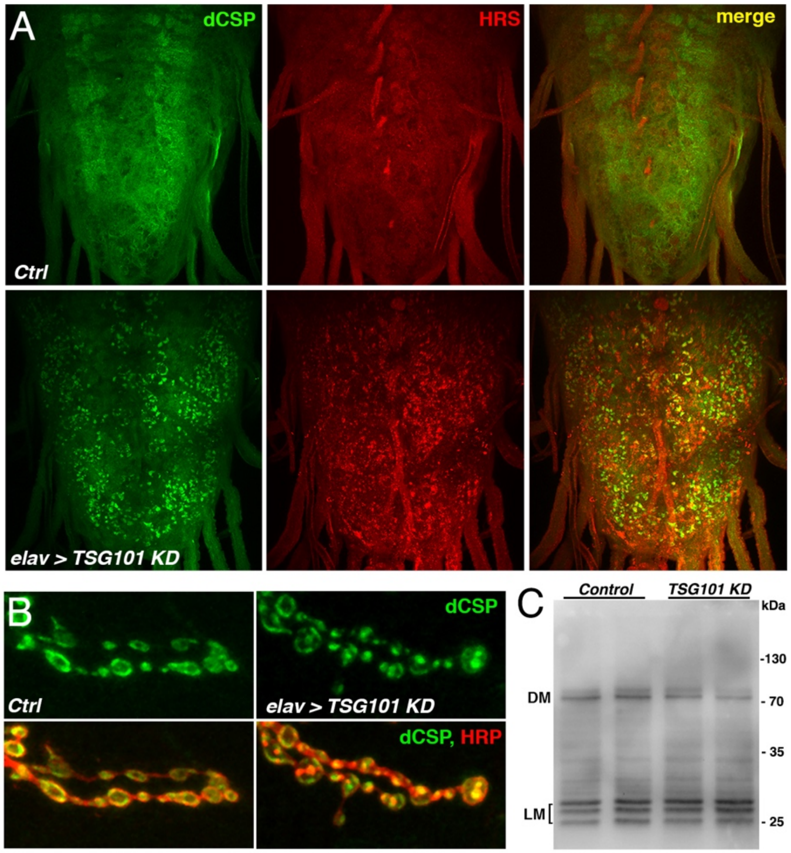


**Figure S4. RNAi-mediated KD of TSG101 causes accumulation of dCSP on HRS-positive endosomes. A.** Larval VNCs immunostained for dCSP and HRS of driverless control (*w*$^{1118}$*; UAS-TSG101 KD/*+) and elav-driven KD of TSG101 using a hairpin transgene (*w*$^{1118}$*, elav-Gal4; UAS-TSG101 KD/*+). Note the mislocalization of endogenous dCSP from its typical diffuse neuropil localization to HRS-positive accumulations in neuropil and neuronal somata. **B.** Larval NMJs of indicated genotypes stained for dCSP and HRP. Note the abnormal localization of dCSP from the periphery to more entrally located HRP-positive endosomes. **C.** Immunoblot probed for dCSP of larval brain extracts from control and elav-driven TSG101 KD. Note that TSG101 KD does not cause high-molecular weight oligomerization of dCSP despite the induced mislocalization to HRS-positive endosomes.

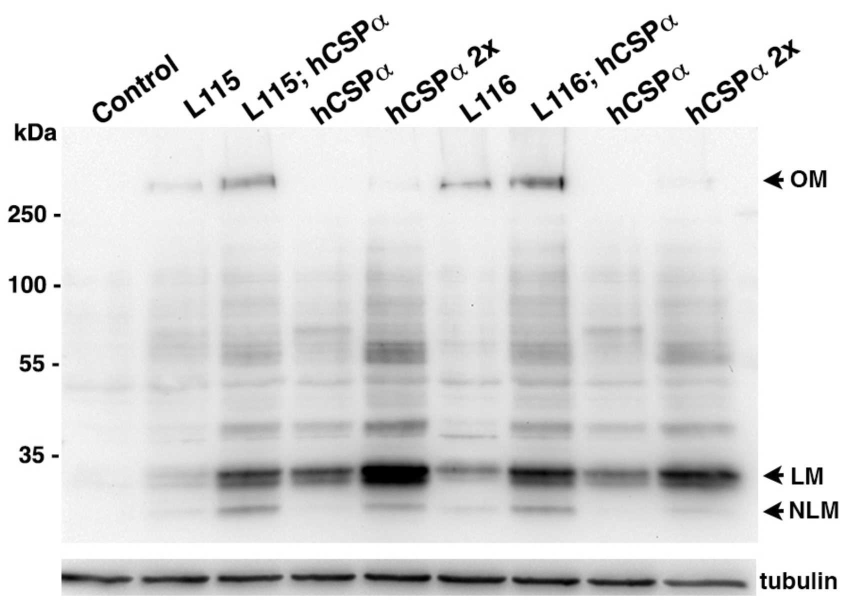


**Figure S5. Effects of dCSP or hCSPα co-expression on hCSP-L115 and -L116 protein levels.** Western blot of larval brain extracts from indicated genotypes probed for hCSPα. WT and mutant hCSPα (L115 and L116) were co-expressed with WT hCSPα or dCSP in larval neurons of control (*$w^{1118}$*) animals with an elav-Gal4 driver. WT hCSPα was also expressed from two transgenes (2x). β-tubulin was used as loading control. hCSPα oligomers (OM), lipidated (LM), non-lipidated and monomers (NLM) are indicated.

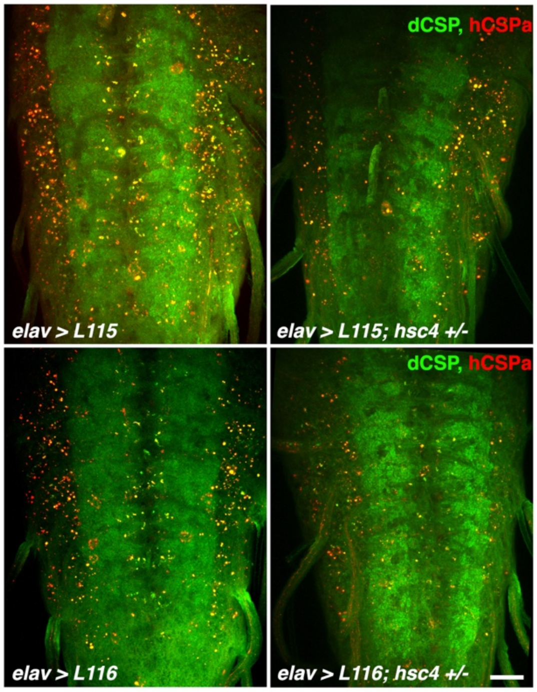


**Figure S6. Effects of altering *hsc4* gene dosage on *CLN4* mutant hCSPα expression.** Larval VNCs for indicated genotypes immunostained for dCSP (green) and hCSPα (red). hCSP-L115 and -L116 were expressed in larval neurons with an elav driver from 1 transgene in wild type control (w*hite*$^{1118}$) or heterozygous *hsc4*$^{\Delta 356}$ deletion mutants. Scale bar, 20 µm.
